# The dynamic epigenetic regulation of the inactive X chromosome in healthy human B cells is dysregulated in lupus patients

**DOI:** 10.1101/2020.11.25.399022

**Authors:** Sarah Pyfrom, Bam Paneru, James J. Knoxx, Michael P. Cancro, Sylvia Posso, Jane H. Buckner, Montserrat C. Anguera

## Abstract

Systemic lupus erythematous (SLE) is a female-predominant disease characterized by autoimmune B cells and pathogenic autoantibody production. Individuals with two or more X chromosomes are at increased risk for SLE, suggesting that X-linked genes contribute to the observed sex-bias of this disease. To normalize X-linked gene expression between sexes, one X in female cells is randomly selected for transcriptional silencing through X-Chromosome Inactivation (XCI), resulting in allele-specific enrichment of epigenetic modifications, including histone methylation and the long noncoding RNA XIST/Xist on the inactive X (Xi). As we have previously shown that epigenetic regulation of the Xi in female lymphocytes from mice is unexpectedly dynamic, we used RNA FISH and immunofluorescence to profile epigenetic features of the Xi at the single cell level in human B cell subsets from pediatric and adult SLE patients and healthy controls. Our data reveal that abnormal XCI maintenance in B cells is a feature of SLE. Using single-cell and bulk cell RNA sequencing datasets, we found that novel X-linked immunity genes escape XCI in specific healthy human B cell subsets, and that human SLE B cells exhibit aberrant expression of X-linked genes and XIST RNA Interactome genes. Our data reveal that mislocalized XIST RNA, coupled with a dramatic reduction in heterochromatic modifications at the Xi in SLE, predispose for aberrant X-linked gene expression from the Xi, thus defining a novel genetic and epigenetic pathway that affects X-linked gene expression in human SLE B cells and likely contributes to the female-bias in SLE.

## INTRODUCTION

Systemic lupus erythematosus (SLE) is an incurable autoimmune disease with multiorgan system manifestations. B cells contribute to various aspects of SLE by secreting pathogenic autoantibodies, presenting autoantigens to T cells, and producing inflammatory cytokines. In addition, the representation of B cell subsets changes in SLE, which can accelerate the production of autoantibodies. In particular, CD27^-^ memory B cells^1^, CD19^hi^CXCR3^hi^ B cells^2^, CD24^-^ -activated naïve B cells^3^, and age-associated CD11c^+^ B cells that express T-bet^4,5^ are increased in autoimmunity. Thus, a further understanding of the mechanisms involved in autoimmune B cell dysregulation is critical for future efforts to control the development and progression of SLE.

Like many autoimmune diseases, SLE exhibits a strong female bias, with 85% of patients being women. The underlying mechanisms responsible for this sex difference are not well understood, yet it is clear that the genetics of the X chromosome impacts disease susceptibility^6^. Indeed, individuals with two or more X chromosomes are at increased risk for SLE^7^, suggesting that X-linked genes have a significant role in disease. Immunity-related genes are enriched on the X chromosome^8,9^, and some of these genes are routinely overexpressed in SLE patient B cells^10-13^. In addition, mouse models with X-linked gene duplication (such as the BXSB-Yaa mouse model^14,15^) or transgenic overexpression of either of the X-linked genes *Tlr7*^16,17^ or *Btk*^18,19^ exhibit disease resembling human SLE, with production of dsDNA autoantibodies. Thus, abnormal dosage or expression of particular X-linked genes is associated with SLE disease in mice and humans.

Female mammalian cells with two X chromosomes regulate X-linked gene expression using X-chromosome Inactivation (XCI), in which one X is randomly selected for transcriptional silencing to equalize gene expression between the sexes^20,21^. Numerous epigenetic modifications, including histone methylation^22,23^, DNA methylation^24,25^, and the long noncoding RNA XIST/Xist^26-28^ are enriched allele-specifically on the inactive X (Xi), and maintain transcriptional repression of most of the X-chromosome. However, some X-linked genes escape XCI, and human cells exhibit higher levels of XCI escape (15-25% of the X chromosome) compared to mice (3% escape)^29,30^. While most somatic cells maintain XCI with static enrichment of Xist RNA and heterochromatin marks on the Xi, we found that lymphocytes exhibit a unique dynamic localization of these modifications to the Xi following stimulation^31-33^. These observations are likely to be significant to pathogenesis, as we recently showed that T cells from SLE patients have dispersed XIST RNA transcripts and aberrant overexpression of many X-linked gene transcripts compared to T cells from healthy controls^32^.

In this study, we determined the epigenetic profile of the Xi in human B cell subsets at the single-cell level, and found that typical heterochromatic modifications are missing from the Xi, suggestive of high levels of XCI escape across B cells. Remarkably, we found mislocalized XIST RNA and reductions with the heterochromatin mark H2AK119Ub at the Xi in activated B cells from pediatric and adult SLE patients, and accordingly, discovered aberrant gene expression profiles of X-linked genes in activated SLE B cells. Our study demonstrates that the unique chromatin features of the Xi in human B cells facilitates XCI escape, and we propose that impaired XCI maintenance in SLE results in aberrant gene expression of X-linked genes, that may further contribute to autoimmunity.

## RESULTS

### Circulating human B cell subsets lack robust XIST RNA localization at the Xi

We previously reported that naïve B cells from female humans lack detectible XIST RNA signals at the Xi, and that naïve B cells from female mice are missing both Xist RNA and enrichment of the heterochromatin modification H3K27me3 at the Xi. XIST RNA localization patterns in lymphocytes can be classified into 4 groups^31,32^: Type I cells have robust XIST RNA localized on the Xi; Type II cells have diffuse XIST RNA signals within a nuclear territory encompassing the X chromosome; Type III cells have XIST RNA pinpoints across the nucleus; and Type IV cells lack XIST RNA signals (Fig. 1A). To determine whether XIST RNA and the heterochromatin modification H2AK119-ubiquitin (H2AK119Ub) were missing from the Xi in human B cell subsets, we isolated circulating naïve B (CD19^+^CD10^−^CD21^+^IgD^+^), memory B (CD19^+^ CD27^+^), plasma (CD19^+^ CD138^+^), and age-associated B cells (ABCs; CD19^+^CD11c^+^) from healthy human donors for sequential XIST RNA FISH and immunofluorescence (IF). Remarkably, we found that both XIST RNA transcripts and H2AK119Ub foci were missing from the Xi in naïve B cells, plasma cells, and ABCs (Fig. 1B). Memory B cells had dispersed XIST RNA signals across the nucleus, yet also lacked H2AK119Ub foci (Fig. 1A). We quantified the XIST RNA localization patterns for human B cell subsets and found that naïve B, ABCs, and plasma B cells are predominantly Type IV, and memory B cells were mostly Type III with some Type II patterns (one-way ANOVA for Type II, Type III, Type IV p < 0.05; Fig. 1C). All four B cell subsets examined lacked detectable H2AK119Ub foci (Fig. 1D), including memory B cells that displayed Type III XIST RNA pinpoints across the nucleus. As a previously published RNAseq dataset^34^ revealed that *XIST* is continuously expressed in naïve, memory, and ABC cells (Fig. 1E), our findings indicate that XIST RNA localization and transcription of the *XIST* gene are genetically uncoupled in human B cell subsets.

**Figure 1:**
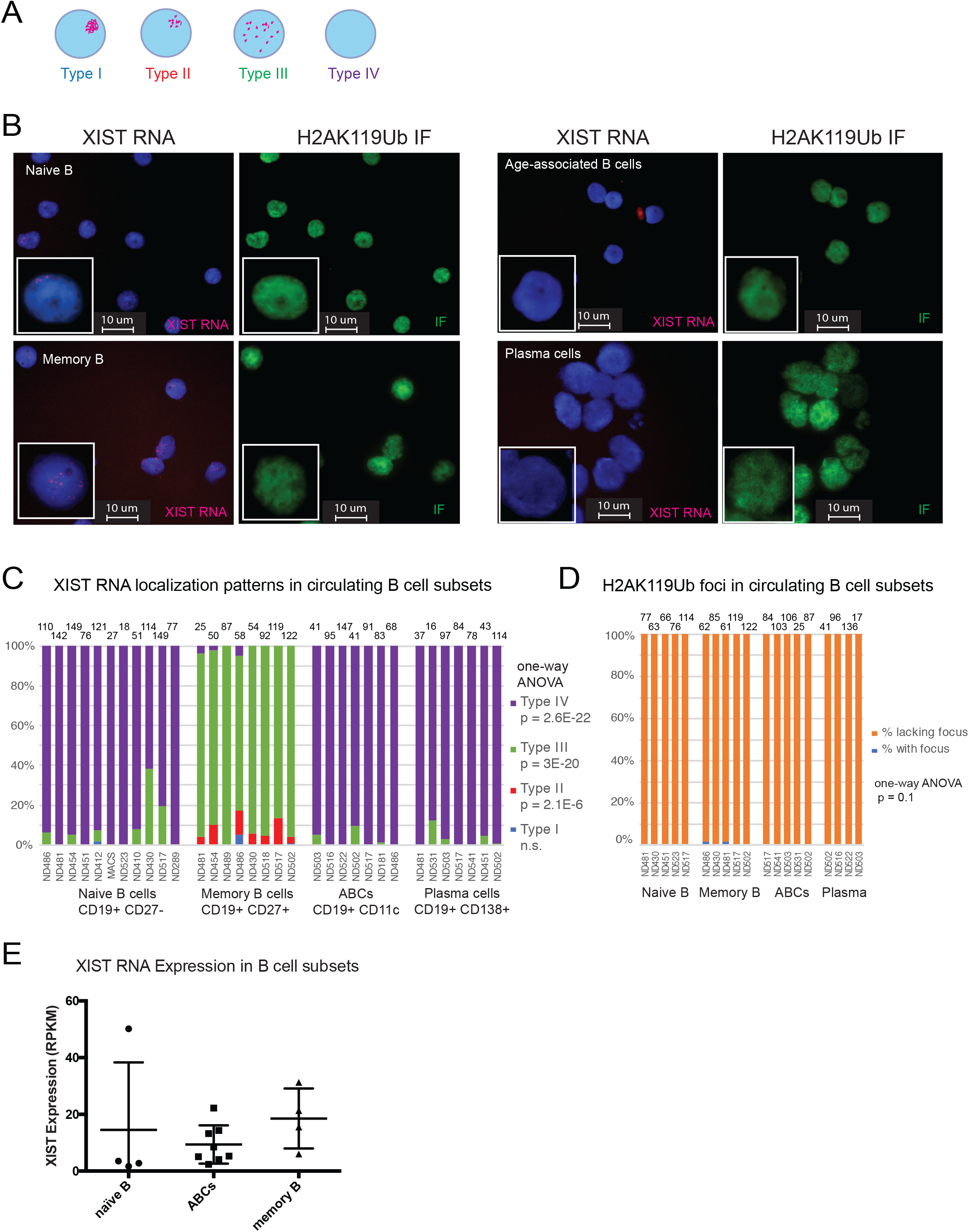
XIST RNA signals are missing from the Xi in human B cell populations. (A) Cartoon representing each type of XIST RNA localization pattern observed in human B cell subsets. (B) Sequential XIST RNA FISH (red) then immunofluorescence detection (green) for H2AK119-ubiquitin (Ub) for naïve B cells, memory B cells, age-associated B cells (ABCs), and plasma cells from healthy PBMCs. (C) Quantification of XIST RNA localization patterns from each B cell subset. Number of nuclei counted is shown above each sample at the top of the graph. Statistical significance for each XIST RNA localization pattern determined using one-way ANOVA; *p* values for each pattern shown. (D) Quantification of H2AK119Ub foci in human B cell subsets. Number of nuclei counted is shown above each sample at the top of the graph. (E) XIST RNA reads for naïve B, ABCs, and memory B cells from a previously published RNAseq dataset^34^.

### XIST RNA and heterochromatin modifications H2AK119Ub and H3K27me3 are localized at the Xi after *in vitro* activation of mature naïve human B cells

While naïve B cells from female mice lack Xist RNA and heterochromatin mark enrichment on the Xi, we have shown that these epigenetic modifications return to the Xi at 24-30 hrs post stimulation^35^. Here, we determined if the dynamic localization of XIST RNA is similarly observed in healthy naïve and memory B cells stimulated *in vitro* using CpG for 3 -7 days. Using RNA FISH, we find that XIST RNA transcripts were first detected in naïve B cells at 1-2 days post-stimulation, and signals decreased by 3-4 days post-stimulation (Fig. 2A). The efficiency of *in vitro* stimulation was assessed by CD86+ staining, with ∼50-60% of B cells being positive for this marker in each XIST RNA experiment (Fig. 2B). We quantified the XIST RNA localization patterns during human B cell activation and found that day 2 stimulated B cells had the highest levels of Type I and Type II XIST RNA localization patterns (Fig. 2C), with such patterns appearing at day 1 post-stimulation. Type III XIST RNA localization patterns were predominant at days 3-7 post *in vitro* activation (Fig. 2C). XIST RNA transcript levels were relatively similar between naïve and *in vitro* stimulated B cells (Supplemental Figure 1), as previously observed in mouse B cells^35^ and reflecting uncoupled XIST transcription and localization to the Xi. Similar analysis of circulating memory B cells revealed that Type I and II XIST RNA patterns predominated after 3 days of culture post-stimulation for memory B cells (Fig. 2D, 2E). In sum, *in vitro* activation using CpG stimulates the return of XIST RNA transcripts to the Xi in both naïve and memory B cells.

**Figure 2:**
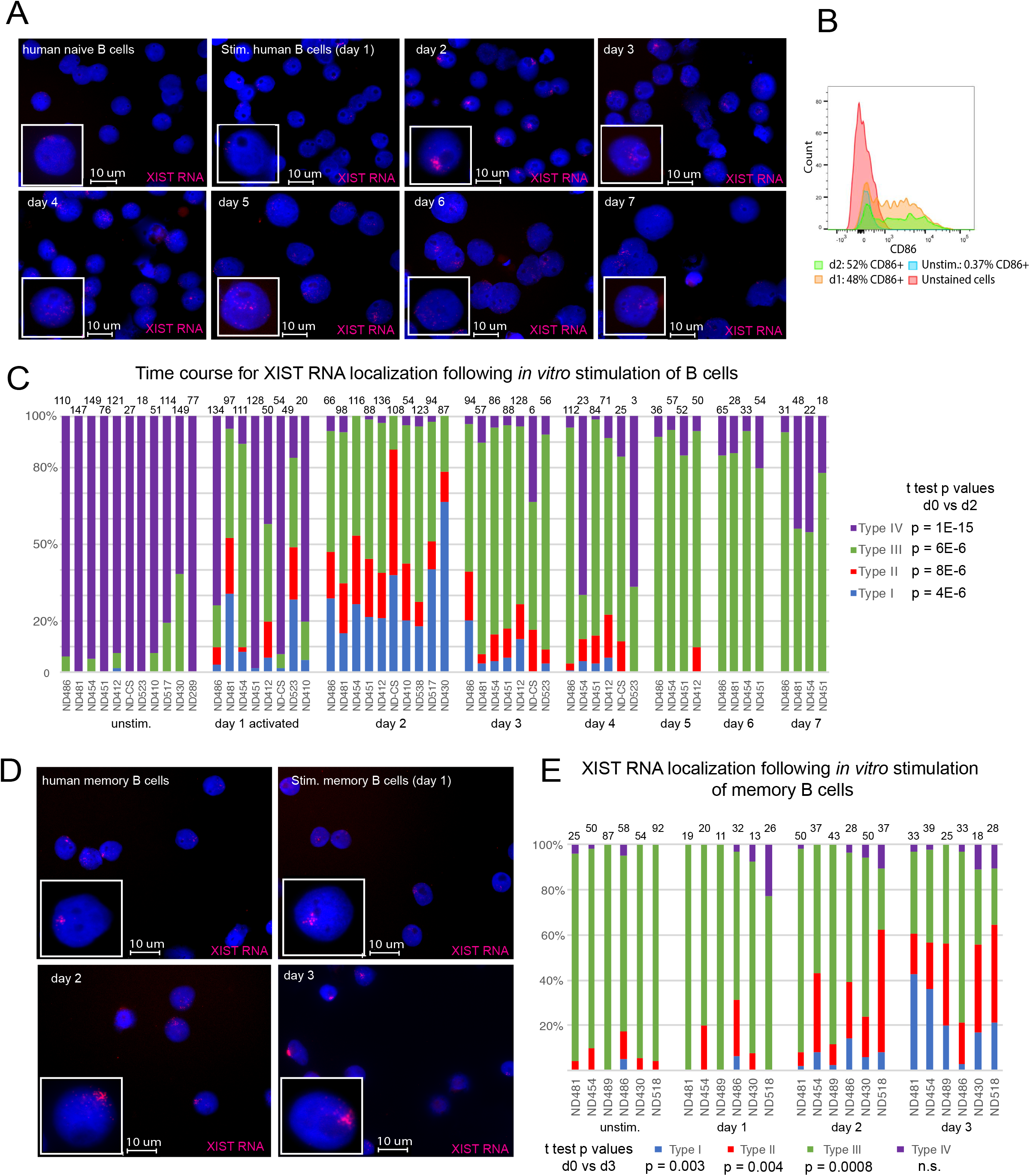
Timing for XIST RNA localization to the Xi during human B cell stimulation, for naïve and memory B cells. (A) Time course analysis for XIST RNA FISH to monitor XIST RNA localization changes after B cell stimulation using CpG, over 7 days. (B) CD86+ staining of in vitro activated B cells (day 1, day 2) to measure efficiency of *in vitro* stimulation. (C) Quantification of XIST RNA localization patterns for *in vitro* stimulated naïve B cells over 7 days in culture. Number of nuclei counted is shown above each sample at the top of the graph. Statistical significance determined using t-test comparing day 0 to day 2, for each type of XIST RNA localization pattern. (D) XIST RNA FISH for *in vitro* stimulated memory B cells using CpG, over 3 days. (E) Quantification of XIST RNA localization patterns for *in vitro* activated memory B cells over 3 days. Number of nuclei counted is shown above each sample at the top of the graph. Statistical significance determined using t-test comparing day 0 to day 3, for each type of XIST RNA localization pattern.

We next asked whether XIST RNA recruitment to the Xi coincided with the enrichment of heterochromatic foci typical of the Xi in somatic cells^22^. Circulating naïve B cells were activated with CpG for 2 days, and then used for sequential XIST RNA FISH followed by IF using antibodies for H3K27me3 and H2AK119Ub. We quantified the number of nuclei that exhibited co-localization of XIST RNA with a heterochromatic focus (Fig. 3). The majority of activated B cells (40-80%) contained a focus that co-localized with Type I XIST RNA patterns and either H2AK119Ub (Fig. 3A) or H3K27me3 (Fig. 3B). There were very few nuclei with XIST Type IV patterns (purple bars), and a focus with either H2AK119Ub or H3K27me3 (2-4%), suggesting that XIST RNA localization to the Xi may be necessary for enrichment of these repressive modifications. Return of epigenetic marks to the Xi during human B cell activation occurs in one phase in which both XIST RNA and heterochromatin modifications appear concurrently at the Xi beginning at day 1 post-stimulation using CpG, with peak enrichment occurring at day 2.

**Figure 3:**
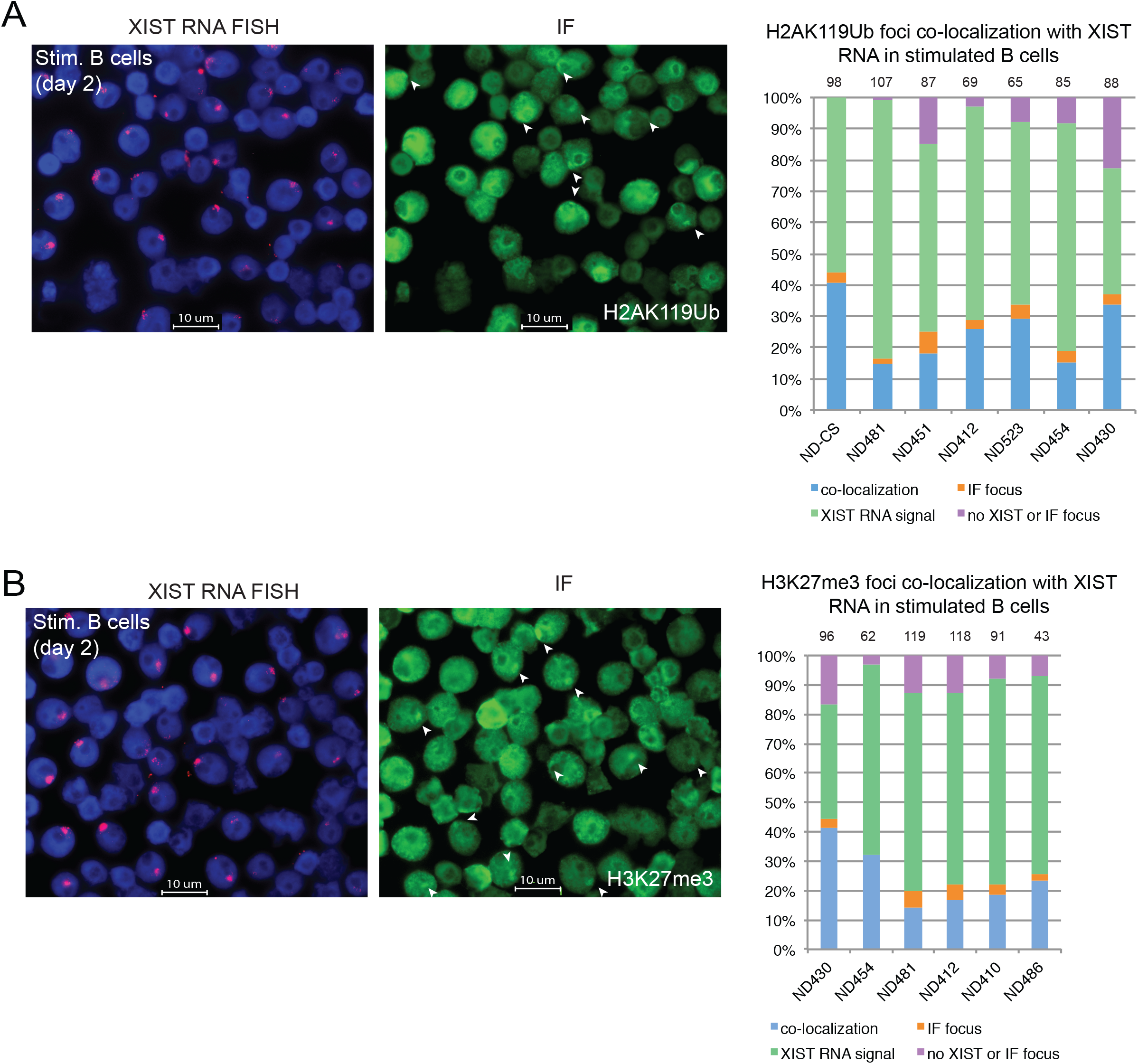
Co-localization of XIST RNA and heterochromatin marks H2AK119Ub and H3K27me for *in vitro* activated human B cells. Sequential XIST RNA FISH (red) followed by immunofluorescence detection (green) for (A) H2AK119Ub and (B) H3K27me3. Representative images (showing the same field) are shown. Quantification of co-localization patterns for XIST RNA and each heterochromatin mark at the Xi. Co-localization of XIST RNA (Types I, II) and IF focus (blue bars), XIST RNA signals alone (Type III; green), nuclei without either signal (purple), or IF focus (orange). Number of nuclei counted is above each sample.

Next, we asked whether the histone variant macroH2A, which is also typically enriched on the Xi in fibroblasts, was localized to the Xi of *in vitro* activated healthy human B cells. We observed very few macroH2A1 foci in these cells, with approximately 10% co-localization of XIST RNA with a focus of macroH2A1 (Supplemental Figure 2A, 2B). Use of qRT-PCR revealed expression of both transcript variants, but these levels still remained below those observed in human female fibroblasts (Supplemental Figure 2C). We also investigated whether the active chromatin modification H3K4me3 was depleted within the territory of the Xi in activated B cells, as typically observed in female fibroblasts^36,37^. Using sequential XIST RNA FISH followed by IF, we observed the characteristic H3K4me3 ‘holes’, reflecting active transcription, which overlapped with XIST RNA Type I and II signals in 70-85% of the nuclei (Supplemental Fig. 3). In sum, the chromatin of the Xi in CpG activated human B cells is enriched for some, but not all, silent and active chromatin modifications, underscoring important differences with other somatic cells.

### Single-cell transcriptional profiling of human B cell subsets reveals cell-type specific biallelic expression of X-linked genes

The absence of XIST RNA and heterochromatic modifications H2AK119Ub, H3K27me3, and macroH2A from the Xi in circulating B cell subsets suggests that there may be either increased overall transcription from this chromosome or that the number of genes that escape from XCI would be increased. To investigate this possibility, we queried single-cell RNA sequencing (scRNAseq) data from a recent study^38^ examining 117 human B cells isolated from a healthy female donor. These cells were sorted by surface markers and consist of 30 memory B cells, 30 naïve B cells, 30 plasmablasts, and 27 transitional B cells. We determined the X-linked SNP expression for genes in each cell using a SNP detection threshold of at least 10 reads/SNP (Fig. 4A). Each X-linked gene containing a SNP was called ‘monoallelic’ if greater than 90% of the reads had the same SNP, otherwise the gene was called ‘biallelic’. We detected 6816 individual X-linked SNPs, and 391 unique X-linked genes that were expressed across the B cell subsets (Fig. 4A; Supplemental Tables 1, 2). We observed novel cell-type specific XCI escape with higher levels of biallelic expression in memory B cells (98 biallelic genes) and plasmablasts (122 biallelic genes) compared to transitional B and naïve B cells (light blue, Fig. 4B; Supplemental Table 3). We detected a total of 190 X-linked genes that escape XCI across all B cell subsets and 77% of these genes were novel XCI escape genes, as they had not been reported previously (Supplemental Table 3). We also observed expression of 53 X-linked immunity-related genes across the B cell subsets, and found that 38 of these genes (72%) were biallelically expressed (light blue, Fig. 4C; Supplemental Table 4). We summarized the expression status for the X-linked immunity-related genes across the four human B cell subsets in Figure 4D. Biallelic expression (light blue) of *DDX3X* in all four B cell subsets was expected, as *DDX3X* ubiquitously escapes XCI in multiple tissues^30^. While *NONO, AP1S2, TSC22D3, AP1S2, IL2RG, SASH3, MSN*, and *CYBB* also escaped XCI across all 4 B cell subsets, the significance of this biallelic expression is unknown as overexpression of these genes has not been reported in autoimmune diseases. As expected, *XIST* was exclusively monoallelic (dark blue) across all B cells (Fig.4D), given its selective expression from the Xi. We also detected variable XCI escape of *TLR7* in naïve B cells and plasmablasts, which has been observed previously by our group and others^31,39^.

**Figure 4:**
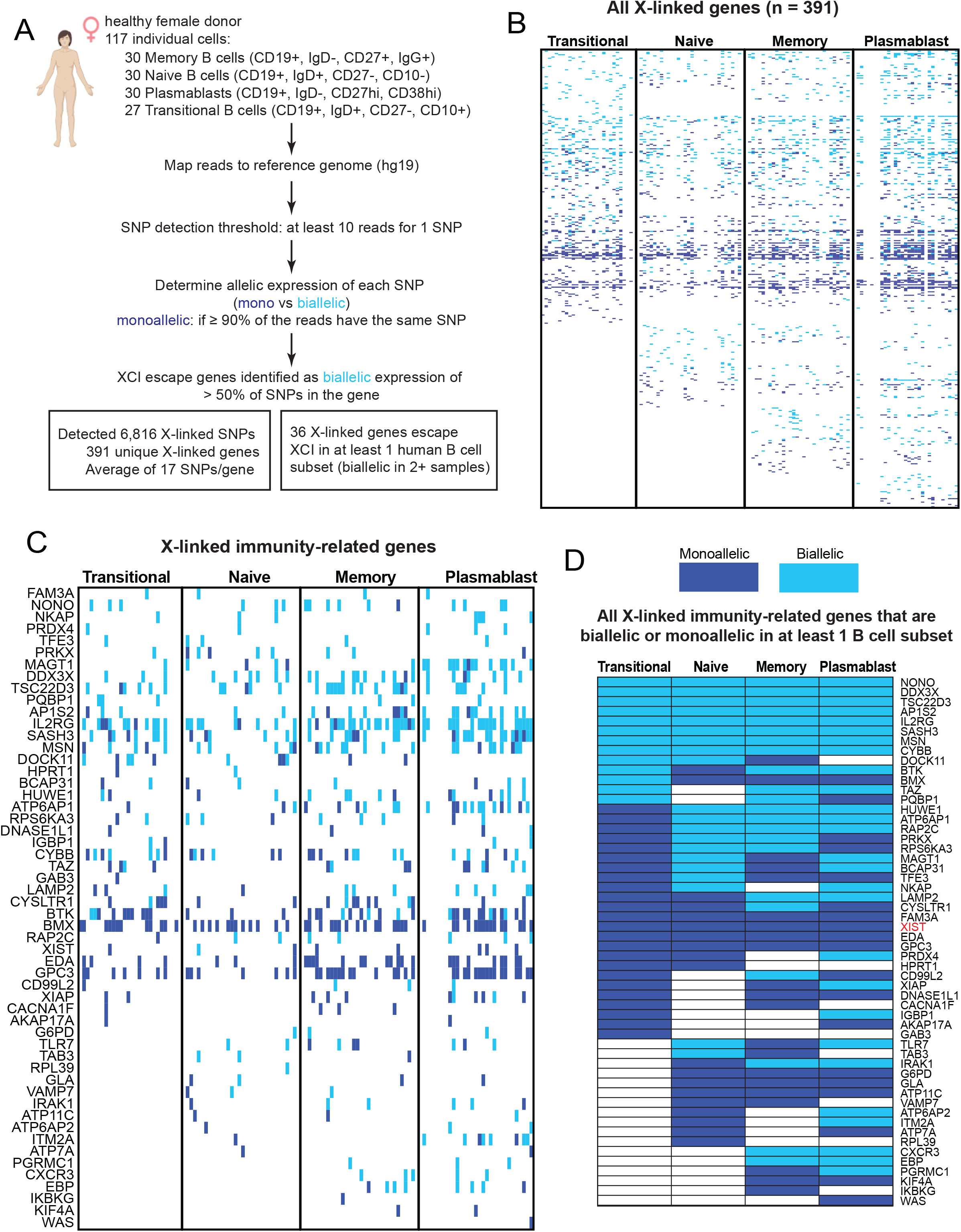
Biallelic expression of X-linked genes in human B cell populations. (A) Schematic of the bioinformatics analysis pipeline to identify XCI escape genes in circulating memory B cells, naïve B cells, transitional B cells, and plamsablasts. (B) Heatmap for all X-linked genes with detectable expression across the four human B cell subsets. Each individual cell is a column. Light blue indicates biallelic expression (Minor Allele Frequency (MAF) ≥ 0.1 for > 50% of SNPs per gene), dark blue indicates monoallelic expression, and white is undetectable expression. Gene lists for individual SNPs in each cell (across B cell populations) are found in Supplemental Table 1, and complete list of all expressed X-linked genes is in Supplemental Table 2. (C) Expression of X-linked immunity-related genes across all four B cell populations. Individual cells for a particular B cell subset shown in columns; monoallelic expression (dark blue); biallelic expression (light blue). Supplemental Table 3 contains gene lists for each B cell subset. (D) Allelic expression summary for the X-linked immunity-related genes, either monoallelic (dark blue) or biallelic (light blue) across human B cell subsets. A gene was considered biallelic for a particular B cell subset if 2 or more cells within that group were biallelic. A gene was considered monoallelic for a B cell subset if 2 or more cells within that subset were monoallelic. Supplemental Table 4 contains complete list of X-linked immunity-related genes that were expressed in each B cell subset, along with allelic expression information.

Interestingly, we also observed variable XCI escape for *BTK*, which is notable because dosage imbalances of this gene are associated with lupus-like phenotypes^18,19^. In sum, human B cells exhibit cell-type specific XCI escape of important immune regulatory genes that could potentially contribute to sex-dependent differences in cellular function, thereby impacting autoimmune disease.

### Pediatric SLE patient B cells have missing or mislocalized XIST RNA transcripts from the Xi and lack H2AK119Ub foci

Cell lines derived from SLE patient B cells exhibited differences in XIST RNA localization patterns compared to cell lines from age-matched, healthy individuals^31^. Here, we investigated whether primary naïve B cells from SLE patients would also have mislocalized XIST RNA patterns. Naïve B cells isolated from pediatric SLE patients in disease remission (SLE disease activity index [SLEDAI] score 0-1) were stimulated *in vitro* using CpG for 2 days, and then used for XIST RNA FISH analyses. We quantified the percentage of each type of XIST RNA localization pattern for both circulating naïve B cells and activated B cells that had been stimulated *in vitro* (Supplemental Figure 4). There were no significant differences between XIST RNA localization patterns for circulating naïve B cells from pediatric SLE patients and age-matched healthy controls (Supplemental Figure 4A). In contrast, we found that there were very few examples of Type I and significantly reduced levels of Type II XIST RNA patterns for pediatric SLE *in vitro* stimulated B cells compared to healthy controls (Fig. 5A, 5B, Supplemental Figure 4B; p < 0.0001, and p = 0.0002). *In vitro* activated SLE patient B cells also had significantly higher levels of Type IV XIST RNA patterns, where nuclei lack detectable XIST signals (Fig. 5B, Supplemental Figure 4B; p = 0.008). Aberrant XIST RNA localization patterns in activated pediatric SLE B cells were not a result of impaired or ineffective stimulation using CpG (a TLR9 agonist), as SLE B cells had similar levels of the activation marker CD86 (Supplemental Figure 4C). XIST RNA localization patterns were also disrupted in SLE patient memory B cells compared to healthy controls, with predominantly Type III and Type IV patterns (Supplemental Figure 4D; p = 0.004). To further assess if enrichment of the heterochromatic modification H2AK119Ub was also affected, we performed IF for H2AK119Ub on circulating and *in vitro* activated B cells from pediatric SLE patients and healthy controls. We observed a significant reduction in H2K119Ub foci in the activated B cells from pediatric SLE patient samples relative to healthy controls (Fig. 5C; p = 8.9E-6). Circulating naïve B cells from both SLE patients and healthy controls lacked detectible H2AK119Ub foci, as expected (Fig. 1). In sum, mislocalization of XIST RNA and near-absence of H2AK119Ub foci on the Xi is a feature of activated pediatric SLE patient B cells in disease remission.

**Figure 5:**
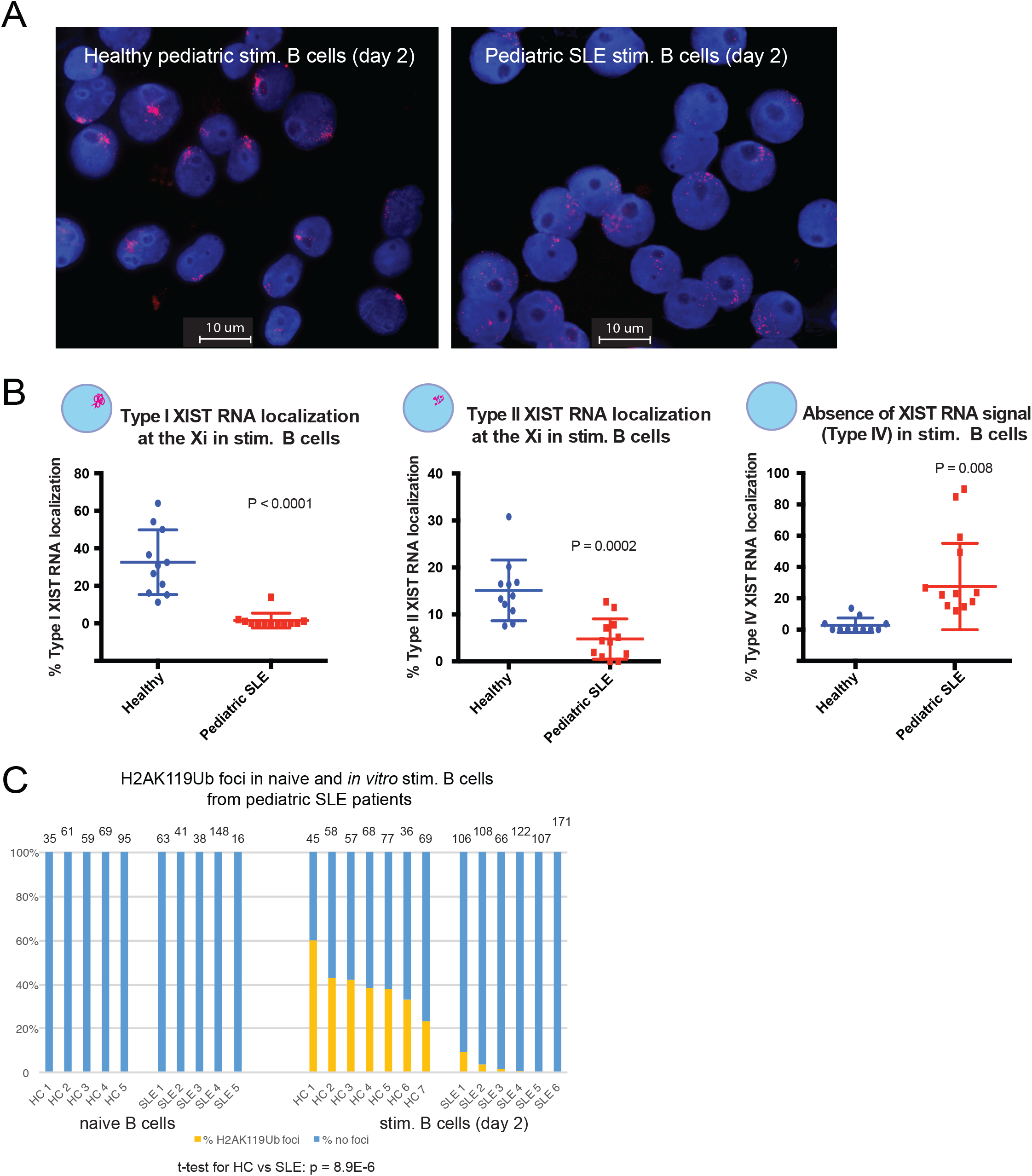
Peripheral B cells from pediatric SLE patients have mislocalized XIST RNA patterns and lack H2AK119Ub foci at the Xi. (A) Representative XIST RNA FISH images from *in vitro* activated B cells (cultured 2 days) from one pediatric SLE patient (right) and a healthy age-matched control (left). (B) Quantification of Type I (left), Type II (center) and Type IV (right) XIST RNA localization patterns for *in vitro* activated B cells from pediatric SLE patients (red) and healthy controls (blue). Error bars denote mean +/-SD, and statistical significance was determined using two-tailed unpaired t-test. (C) Quantification of H2AK119Ub foci for *in vitro* activated B cells from pediatric SLE patients and healthy control samples. Number of nuclei counted is above each sample; statistical significance comparing SLE to healthy controls was determined using two-tailed unpaired t-test. (D) Quantification of XIST RNA localization patterns for *in vitro* activated classical memory B cells, cultured for 3 days with CpG. Number of nuclei counted is above each sample; statistical significance comparing SLE to healthy controls was determined using two-tailed unpaired t-test for each pattern of XIST RNA localization.

### B cells from adult SLE patients have aberrant XIST RNA localization patterns and significant reductions in H2AK119Ub enrichment on Xi, irrespective of disease activity

We initially confirmed that that CD27-isolated cells from SLE patients had relatively similar representation of all B cell populations relative to healthy controls (Supplemental Figure S5). Similar to *in vitro* activated pediatric SLE patient B cells, *in vitro* activated B cells from adult SLE patients had significantly fewer Type I (p < 0.0001) and Type II (p < 0.0001) XIST RNA patterns, and significantly more Type IV patterns (p < 0.0001), signifying abnormal XIST RNA localization patterns in B cells from adult SLE patients (Figures 6A, 6B; Supplemental Figure 6). In contrast to the pediatric SLE population, about half of the adult SLE patients had similar levels of Type I and Type IV XIST RNA patterns as healthy controls (Fig. 6B). Co-localization of XIST RNA with H2AK119Ub foci were significantly reduced in adult SLE patient B cells compared to healthy controls (blue bars, p < 0.0001; Fig. 6C), and cells containing an H2AK119Ub focus independent of XIST RNA signals (Types I, II) were absent in SLE samples (orange bars). Activation of naïve B cells following stimulation with CpG (as determined by CD86+ levels) was not significantly affected in adult SLE samples (Supplemental Figure 6B). In sum, enrichment of both XIST RNA and the heterochromatic modification H2AK119Ub at the Xi are significantly reduced for *in vitro* activated adult SLE B cells.

**Figure 6:**
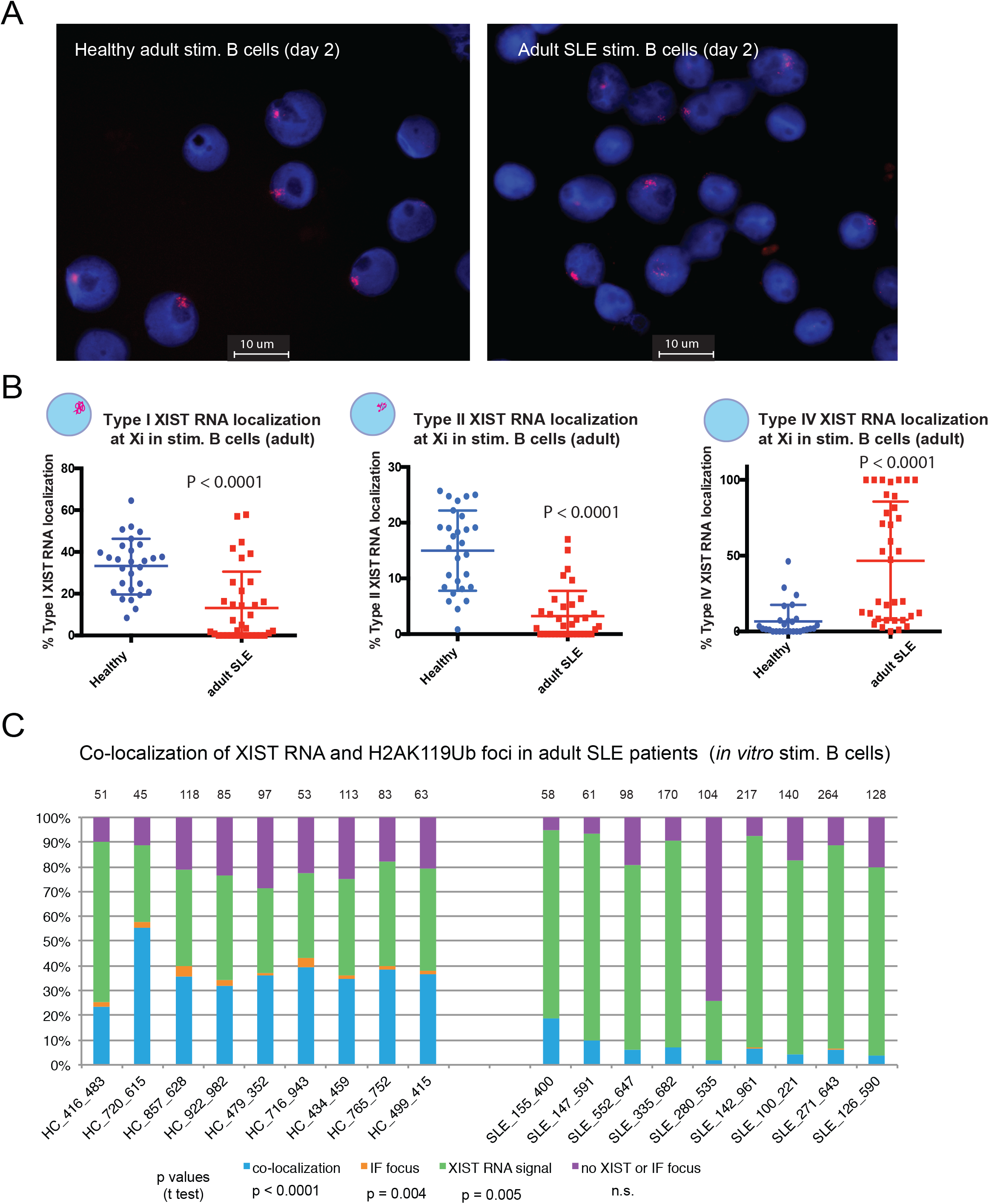
Peripheral B cells from adult SLE patients have mislocalized XIST RNA patterns and reduced H2AK119Ub foci at the Xi. (A) Representative XIST RNA FISH images from *in vitro* activated B cells (cultured 2 days) from one adult SLE patient (right) and a healthy age-matched control (left). (B) Quantification of Type I (left), Type II (center) and Type IV (right) XIST RNA localization patterns for *in vitro* activated B cells from adult SLE patients (red) and healthy controls (blue). Error bars denote mean +/-SD, and statistical significance was determined using two-tailed unpaired t-test. (C) Quantification of co-localization patterns for XIST RNA and H2AK119Ub at the Xi, for *in vitro* activated B cells from adult SLE patients and healthy control samples. Co-localization of XIST RNA (Types I, II) and IF focus (blue bars), XIST RNA signals alone (Type III; green), nuclei without either signal (purple), or IF focus (orange). Number of nuclei counted is above each sample; statistical significance comparing SLE to healthy controls was determined using two-tailed unpaired t-test for each pattern of XIST RNA localization.

We next asked whether the distribution of XIST RNA patterns correlated with SLE disease activity (SLEDAI values), patient medications, age, or disease duration. For these analyses, we removed patients with known comorbidities, including thyroid illnesses (such as Grave’s disease) as they exhibit greater Type III and fewer Type IV XIST RNA patterns (Supplemental Figure S7; left panels). Analysis of each XIST RNA localization pattern and the 6 medications typically used to treat SLE symptoms demonstrated that only hydroxychloroquine showed significant correlation with Type I XIST RNA localization patterns, comparable to healthy controls (Supplemental Fig. 7A,7E). We then performed multiple linear regressions between continuous patient metrics (age at sample draw, SLEDAI score, anti-nuclear antibody titer, and disease duration) and the percentage of Type I-IV XIST RNA localization patterns (Supplemental Figure S8). Of all supplied metrics, SLE patient age was the only one that correlated positively with the Type IV XIST RNA patterns (*p*-value: 0.030, R^2^: 0.214; Supplemental Fig. S8B). Healthy control samples did not exhibit a similar correlation (Supplemental Fig. S8B). Taken together, activated B cells from adult SLE patients exhibit aberrant XIST RNA localization and reduced H2AK119Ub enrichment on the Xi, irrespective of disease activity but correlated with patient age, suggestive of impairments with gene expression on this chromosome.

**Figure 7:**
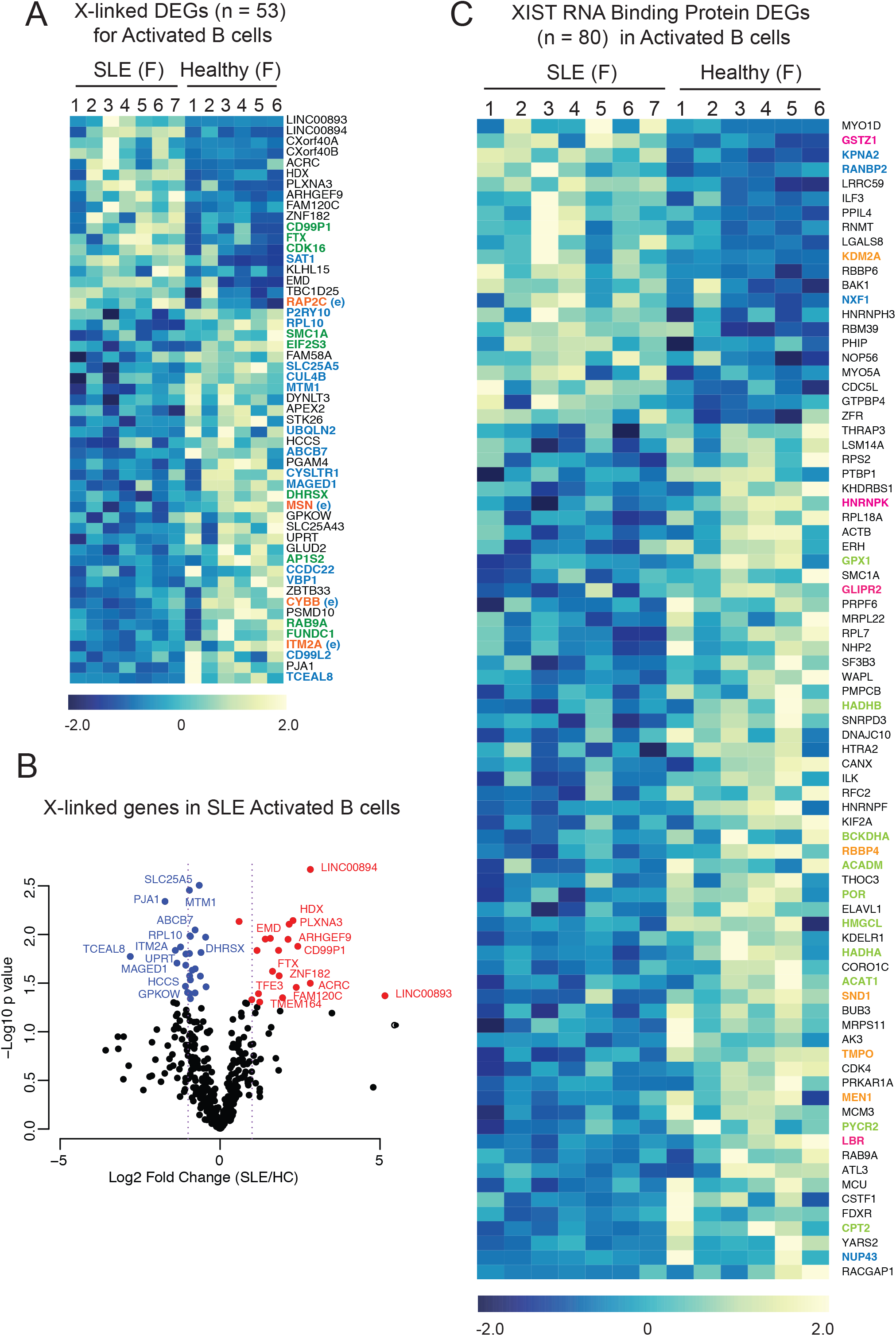
X-linked gene expression and XIST RNA Interactome genes are altered in SLE patient activated B cells. (A) X-linked Differentially Expressed Genes (DEGs; 53 genes) in adult healthy controls (6 female samples) and SLE patient (7 female samples) activated B cells in circulation. Color gradient represents row Z-scores for each gene. Gene symbols in color denote XCI escape: orange are immunity-related genes that may escape in activated B cells; green are known XCI escape genes in other somatic cells; blue are putative XCI escape based on Supplemental Table 3. Note that all 4 genes in orange exhibited XCI escape in at least 1 B cell subset in Supplemental Table 3, and are denoted with (e). (B) Volcano plot showing X-linked DEGs in activated B cells. Genes significantly upregulated in SLE patients are in red; genes significantly downregulated in SLE are in blue (p < 0.05). (C) XIST RNA binding protein genes that are differentially expressed in activated B cells from SLE patients and healthy controls. Nuclear matrix/nuclear envelop genes in blue; cell metabolism/cell growth genes in green; chromatin regulators in orange; XIST RNA binding protein genes whose expression was also altered in SLE patient T cells in pink.

### Activated B cells from SLE patients exhibit abnormal expression of X-linked genes and XIST RNA Interactome genes

Because SLE activated B cells from both pediatric and adult SLE patients exhibited abnormal XIST RNA localization and reduced/missing H2AK119Ub enrichment at the Xi, we asked whether these changes were also associated with abnormal X-linked gene expression. To answer this, we utilized a previously published RNA sequencing dataset (GSE118254) that profiled activated B cells in circulation (CD19+IgD+CD27-MTG+CD24-CD38-) in seven female SLE patients and six healthy controls^40^. In this dataset, we found that 53 X-linked genes were differentially expressed in activated B cells from SLE patients, and 18 of these genes were overexpressed (Figure 7A). Among the 53 differentially expressed X-linked genes, 4 have known immune functions (genes in orange), and these genes exhibited XCI escape in at least 1 B cell subset (Fig. 4D, Supplemental Table S3). Comparison of the 53 X-linked genes altered in SLE with our results from Supplemental Table 3 and published lists of XCI escape genes from various cell types^30^ indicate that the majority of the genes in Fig. 7A (all the genes in color) may escape XCI in activated B cells. Notably, 18 X-linked genes were significantly upregulated in SLE patient activated B cell samples (red genes), and few of the downregulated genes (blue genes) passed the significance threshold (Fig. 7B). Unexpectedly, we found that the majority of these putative XCI escape genes are downregulated in SLE B cells (Fig. 7A), suggesting impairments with the regulation of XCI escape on the Xi in SLE. In sum, we identified a novel set of X-linked genes whose expression is altered in SLE patient activated B cells, and the majority of these genes should be subject to XCI silencing, reflecting aberrant gene regulation on both Xs.

One possible mechanism for aberrant XIST RNA localization in SLE patient activated B cells could result from impairments with nuclear proteins that bind XIST RNA. The XIST RNA Interactome consists of ∼275 proteins^41-44^, and we have previously reported that two XIST RNA binding proteins, YY1 and hnRNP-U, are required for localization of XIST RNA and heterochromatin marks to the Xi in lymphocytes^31,35^. Notably, SLE patient T cells have altered expression of XIST RNA Interactome genes^32^. Thus, we asked whether the expression of genes encoding XIST RNA binding proteins was also abnormal in the activated B cells from SLE patients. We found that 80 XIST RNA binding protein genes were differentially expressed in activated B cells, and the majority of these genes (59/80; 74%) were downregulated (Fig. 7C, Supplemental Table 5). These genes function in cell metabolism/cell growth (genes in green), nuclear matrix/nuclear envelope/transport (blue, but also includes *LBR* and *hnRNPK*), and chromatin regulation (orange) (Fig. 7C). Downregulated genes *LBR, hnRNP K*, and *GLIPR2* (pink) also exhibited altered gene expression among SLE T cells relative to healthy controls^32^. Taken together, the XIST RNA Interactome is dysregulated in activated B cells from SLE patients, and may be responsible for mislocalization of XIST RNA and heterochromatin modifications from the Xi, resulting in aberrant XCI maintenance.

## DISCUSSION

B cells contribute to the pathogenesis of SLE, an autoimmune disease that predominantly affects women. Here we sought to investigate the genetic basis for the female-bias of SLE, focusing first on how XCI is maintained across distinct B cell subsets in healthy individuals, and then determining if XCI maintenance and X-linked genes expression are affected in SLE. We discovered a diverse enrichment of epigenetic modifications at the Xi across activated human B cell subsets, and found B cell-specific patterns of XCI escape in healthy adults, including the escape of important immune-related genes. Our profiling of pediatric and adult SLE patient B cells revealed significant impairments with XIST RNA and H2AK119Ub enrichment on the Xi irrespective of disease activity, and aberrant expression of X-linked genes. Together, our results suggest that facultative chromatin of the Xi is relaxed in healthy B cells, thereby altering XCI maintenance and permitting gene-specific escape from XCI. SLE disease further impacts the heterochromatic composition of this chromosome, resulting in abnormal gene expression changes across the X. Our epigenetic profiling of the Xi in healthy and SLE B cells provides a foundation for future studies investigating the molecular mechanisms of XIST RNA and heterochromatin mark localization and spreading across the Xi, and how these mechanisms become altered in SLE.

All four human B cell subsets assessed – naïve B, classical memory B, plasma cells, and ABCs – are missing XIST RNA and the heterochromatic modification H2AK119Ub on the Xi (Figure 1). Unexpectedly, our analyses show that unlike naïve B cells, ABCs, and plasma cells, memory B cells have XIST RNA transcripts dispersed across the nucleus, yet still lack H2AK119Ub foci on the Xi (Figure 1C, 1D). At present, the significance of dispersed XIST RNA transcripts in memory B cells is unknown. However, it is likely to impact gene expression on the Xi as *ex vivo YY1* deletion generates similar Type III dispersed patterns with altered expression of ∼70 X-linked genes^35^. The *XIST* gene is expressed across all resting human B cell subsets (Figure 1E), thus transcriptional changes do not account for the absence of XIST RNA transcripts on the Xi in circulating human B cells. XIST RNA and the heterochromatin modifications H2AK119Ub, H3K27me3, and low levels of macroH2A returned to the Xi when using CpG to activate human naïve B cells (Figure 2, 3, Supplemental S2). As for mouse naïve B cells, XIST RNA and heterochromatin marks return to the Xi in human naïve B cells before the first cell division^35^, with peak enrichment at day 2 post-stimulation and prior to cell division^45^. We propose the Xi chromatin in circulating human B cell subsets is more relaxed compared to somatic cells, in which XIST RNA and heterochromatin marks are localized to the Xi, and this may allow additional X-linked genes to escape transcriptional silencing. However, as our cytogenetic RNA FISH and IF analyses lack resolution at the gene-level, it will be important to determine allele-specific enrichment of silent and active histone modifications at genes exhibiting cell-specific XCI escape and silencing across B cell subsets, and to further assess if such modifications are altered upon activation.

Single-cell RNAseq profiling of four human B cell subsets from a healthy female individual reveals cell-type specific XCI escape in naïve B cells, memory B cells, plasmablasts, and transitional B cells. The percentage of biallelically expressed X-linked genes increase with B cell differentiation (Supplemental Table 3), which may reflect a requirement for higher dosage of X-linked genes for proper function of memory B cells and plasmablasts. For example, *LAMP2*, a lysosomal protein important for autophagy and intracellular antigen presentation, is monoallelically expressed in transitional and naïve B cells, yet biallelically expressed in memory B and plasmablasts (Fig. 4D). Such results raise the intriguing possibility that biallelic expression of *LAMP2* may increase autophagy and antigen presentation, thereby contributing to enhanced immune responses observed in females^46^. The X-linked gene *IRAK1*, responsible for IL1-induced upregulation of NF-kappa B, is also biallelically expressed in memory B cells and plasmablasts (Fig. 4D). Female neonates have higher levels of *IRAK1* mRNA and protein in cord blood and mononucleated cells compared to males^47^, potentially contributing to reduced infection rates and female-specific immune advantages in infants. However, these scRNAseq analyses are limited by the fact that the sample was taken from one individual and does not contain *in vivo* stimulated B cells, which may have distinct XCI escape profiles. As XCI escape exhibits individual variability when comparing across human samples^48^, it will be important to repeat the allelic expression profiling for human B cell subsets, especially activated B cells, using healthy female individuals of different ages to determine which X-linked genes consistently escape transcriptional silencing, and whether XCI escape increases with age.

Our investigations revealed abnormal XCI maintenance, as evidenced by reduced XIST RNA and H2AK119Ub enrichment at the Xi, in both pediatric and adult SLE patient B cells, irrespective of disease activity. Perturbed XIST RNA and heterochromatin mark localization to the Xi in lymphocytes is a feature of both human SLE and the analogous lupus-like disease in the female-biased spontaneous mouse model NZB/W F1^32,49^. Activated SLE patient B cells exhibit altered expression of about 50 X-linked genes (Fig. 7A), and the majority of these genes were downregulated in SLE patients compared to healthy controls. It is surprising that more than half of the downregulated X-linked genes in SLE samples are putative XCI escape genes (Fig. 4 and previous studies in human fibroblasts). The implications of reduced expression of these X-linked genes for B cell function is unknown at this time. It is possible that aberrant XIST RNA localization and reduced heterochromatic enrichment reflects alterations in the nuclear organization of the Xi in SLE patient B cells. The Xi, unlike the active X and autosomes, is organized into two “megadomains” separated by a boundary region near the microsatellite repeat *Dxz4*^50,51^. During XCI initiation, Xist RNA plays an important structural role for configuring the Xi territory, and *Xist* deletion impairs megadomain formation^52^. XIST RNA mislocalization in SLE patient B cells may reflect impairments to the Xi nuclear territory, possibly resulting in abnormal gene silencing of some X-linked genes.

While altered X-linked gene expression can clearly impact SLE progression, our studies cannot determine if abnormal XCI maintenance causes SLE disease, or is instead a consequence of the disease. To date, there is no evidence that changes to the extracellular environment can influence XCI maintenance in somatic cells. However, the nuclear pore complex *NUP43* and lamin B receptor (*LBR)*, which are Xist RNA binding proteins, were downregulated in SLE patient B cells (Fig. 7C). This may contribute to aberrant organization of the Xi nuclear territory in SLE patient B cells, as LBR protein directly binds Xist RNA and this interaction is necessary for tethering the Xi to the nuclear lamina and gene silencing^53^. While it is currently unclear if the inflammatory environment of SLE affects nuclear architecture, it is tempting to speculate that inflammatory cytokines or type I interferons, which are highly elevated in SLE patients experiencing disease flares, may perturb XCI maintenance in B cells, resulting in altered X-linked gene expression. Future studies to determine whether extrinsic factors can influence XCI maintenance in lymphocytes will certainly reveal exciting new insights into genetic and epigenetic factors responsible for sex-biased autoimmune disease.

## METHODS

### Human B cell samples from healthy donors and SLE patients

Fresh and frozen PBMCs from adult healthy female donors were obtained from the Penn Pathology BioResource core facility at the Perelman School of Medicine, University of Pennsylvania. For comparative study of pediatric SLE patients (SLEDAI score = 0) and age-matched healthy controls, we recruited patients from the Children’s Hospital of Philadelphia (CHOP). Approximately 15-20 mL of blood were collected from each individual, stored on ice, then immediately processed for PBMC isolation. PBMCs were separated from whole blood by density gradient centrifugation technique using Lymphoprep media (cat # 07851, STEMCELL technologies, Cambridge, MA, USA). PBMCs from CHOP patients were either frozen or used to isolate B cells immediately, and we did not observe any effect of freeze/thaw on XIST RNA localization patterns. PBMCs were frozen in fetal bovine serum containing 7%-10% DMSO. We also obtained frozen PBMC samples from adult SLE patients (SLEDAI score: 0-20; age 18-63 yrs.) and age-matched healthy controls from the Benaroya Research Institute, Seattle, Washington. The acquisition of blood samples from pediatric SLE patients and healthy controls from CHOP was approved by the IRB at CHOP; acquisition of blood from adult SLE patients and healthy controls from the Benaroya Institute was approved by the IRB at the Benaroya Institute. Written informed consent was received from participants prior to inclusion in both studies.

### Sorting and culture of human B cells

Frozen PBMCs were quickly thawed and washed twice with RPMI media containing FBS. CD19^+^CD27^-^ naïve B cells and CD19^+^CD27^+^ memory B cells were isolated from PBMCs using the Easysep human memory B cell isolation kit, according to the manufacture’s instruction (17864, STEMCELL Technologies, Cambridge, MA, USA). CD19^+^CD138^+^ plasma cells were isolated from PBMCs by positive selection using biotin conjugated CD138 antibody (352322, Biolegend, San Diego, CA, USA) and Easysep release human biotin positive selection cocktail (17653, STEMCELL Technologies). CD19^+^CD11c^+^ B cells (ABCs) were isolated from PBMCs using a two-step procedure. First, total B cells were isolated from PBMCs by negative selection using the human B Cell Isolation Kit II (130-091-151, Miltenyi Biotech, Cambridge, MA, USA). ABCs were isolated from purified total B cells by CD11c positive selection using a biotin conjugated CD11c antibody (301612, Biolegend) and anti-biotin magnetic beads. B cells were cultured in X-VIVO™15 media (04-744Q, Lonza, Walkersville, MD, USA) with penicillin-streptomycin (100 units/mL) and activated using 3 µM CpG (ODN 7909) (tlrl-2006-1, Invivogen, San Diego, CA, USA). Cells were cultured in 200 µl medium for 1-8 days using round bottom 96-well plates. B cell stimulation was determined by staining for the activation marker CD86, which was quantified using flow cytometry.

### Flow cytometry profiling of naïve B cells (CD19^+^CD27^-^) from SLE and HC subjects

We used multicolor flow cytometry analysis to determine the subset distribution B cells in SLE and HC samples^54,55^. B cell subsets were phenotyped as follows: naïve B cells (CD19^+^CD10^−^CD21^+^IgD^+^), transitional B cells (CD19^+^CD10^+^CD38^+^), memory B cells (CD19^+^CD27^+^), plasma B cells (CD19+CD138+), and ABCs (CD19^+^CD10^−^CD21^-^ CD85j^+^). Antibodies (and catalog numbers) used for flow cytometry were: FITC CD85j (555942, BD Biosciences), Brilliant Violet 421™ CD38 (303525, Biolegend), Brilliant Violet 650™ CD27 (302827, Biolegend), Brilliant Violet 785™ CD19 (302239, Biolegend), APC/Cy7 CD3 (300425, Biolegend), APC/Cy7 CD14 (561709, BD Biosciences), APC/Cy7 CD16(561726, BD Biosciences), PE-CF594 IgD (562540, BD Biosciences), PE/Cy7 CD21 (354911, Biolegend), BV605 CD24 (311123, Biolegend), PECy5 CD10 (15-0106-41, Thermo Fisher) and LIVE/DEAD™ Fixable Aqua Dead Cell Stain Kit (L34965, Thermo Fisher).

### Sequential XIST RNA FISH and immunofluorescence (IF)

Sequential XIST RNA fluorescence in situ hybridization (FISH) and IF was performed using established protocols ^31,32^. Briefly, cells were cytospun onto glass slides, then incubated in ice-cold cytoskeletal (CSK) buffer containing 0.5% Triton for 3 min, fixed in 4% paraformaldehyde for 10 min, and then dehydrated using an ethanol series. For human XIST RNA FISH, we used two Cy3 labelled oligonucleotide probes which target repetitive regions within *XIST* exons 1, 3 and 4 ^31^. Images were obtained using a Nikon Eclipse microscope and were categorized by the type of XIST RNA localization patterns as shown in Figure 1B and as described previously^31,33,35^. For IF analyses, slides were blocked for 30 min in blocking buffer (PBS with 0.2% Tween-20 and 5% BSA) and then incubated for 2 hours at room temperature with respective primary antibodies (at dilutions of 1:100): H3K27me3 (39155, Active Motif); Ubiquityl-histone H2A Lys119 (8240, Cell Signaling); H3K4me3 (ab 213224, Abcam); MacroH2A1 (ab37264, Abcam). Slides were incubated with the appropriate FITC conjugated secondary antibody for 1 hour at room temperature, then imaged using a fluorescence microscope.

### quantitative RT-PCR

For RT-QPCR, cDNA was synthesized from 1 µg of total RNA using Verso cDNA synthesis kit (Ref # AB-1453/A, Thermo Fisher, Waltham, MA, USA) and qPCR was performed using the QuantBio low ROX cyber green master mix (# 017707, Quantabio, Beverly, MA, USA). RPL13A gene was used as an endogenous control to normalize gene expression. RT-QPCR data were analyzed using ΔΔCt method^5^. RPL13A, F: GCCATCGTGGCTAAACAGGTA, R: GTTGGTGTTCATCCGCTTGC; macroH2A1.1, F: GGCTTCACAGTCCTCTCCAC, R: GGTGAACGACAGCATCACTG; macroH2A1.2, F: GGCTTCACAGTCCTCTCCAC, R: GGATTGATTATGGCCTCCAC, XIST 5’ end: F: TTGCCCTACTAGCTCCTCGGAC, R: TTCTCCAGATAGCTGGCAACC; XIST 3’ end: F: CTACAAGCAGTGCAGAGAGC, R: CTAAGACAAGACACAGACCAC.

### RNA-seq analysis of activated B cells from SLE patients and controls

The processed gene expression file “GSE118254_SLE.RNAseq.geneRpkm.detected.RPM.3.exon.csv.gz” was downloaded from GEO dataset GSE118254 (https://www.ncbi.nlm.nih.gov/geo/query/acc.cgi?acc=GSE118254). Statistical significance in gene expression between Healthy and SLE B cells was calculated using a one-way ANOVA in R (alpha = 0.05). A Z score was calculated for each sample/gene and heatmaps were generated using the gplots (heatmap.2) R package.

### Allele specific gene expression analyses using single cell RNAseq data

Single cell RNAseq reads sequenced from individual naïve B cells, memory B cells, plasmablasts and transitional B cells were retrieved from a previously published study (Bioproject accession number PRJEB27270)^38^. In this study, a total of 117 cells, consisting of 30 naive B cells, 30 memory B cells, 30 plasmablasts and 27 transitional B cells, were sequenced from a single healthy woman. Public servers available through the Galaxy web platform (*usegalaxy*.*org*) were used to analyze the data^56^. RNA-Seq reads were mapped to human reference genome (version hg19) using Bowtie^57,58^. Next, SNPs present in both X chromosomes were detected from alignment file using the FreeBayes variant detector tool (www.geneious.com)^59^ and SNPs were annotated using the ANNOVAR tool^60^. SNPs with less than 10 reads were excluded from the downstream analyses. For each SNP, if ≤ 90% of the reads carry the same SNP allele, expression was considered biallelic, otherwise expression was considered monoallelic for each gene. A gene was considered ‘biallelic’ in a cell-type subset if 2 or more cells from that subset were considered ‘biallelic’ as described above; a gene was considered “monoallelic” is 2 or more cells were biallelic and “uncalled” if no two cells were in agreement or there was insufficient data to make a call.

### Statistical analysis

Linear regression was performed using a simple linear regression in GraphPad Prism 8 of dependent variables (Xist RNA Cloud Type I-V) and independent variables (Date of blood draw for PBMC sample, SLE disease duration, Anti-Nuclear Antibody Titer, Systemic Lupus Erythematosus Disease Activity Index (SLEDAI) score and patient age at blood draw). Measurements of percent Type I-IV XIST RNA clouds were analyzed using unpaired, two-tail t-tests, where significance is p < 0.05.

### Study approval

For pediatric SLE patients, protocols and informed consent forms were approved by the CHOP Institutional Review Board (CHOP IRB Protocol 14-011433). For the adult SLE patients, the studies were approved by the Benaroya Research Institute Institutional Review Committee (IRB protocol number: 10059).

## Figure Legends

**Supplemental Figure 1: Steady-state XIST RNA transcript levels in human naïve and *in vitro* stimulated B cells**. Primer sets for 5’ XIST (spanning exons 1 and 3) and 3’ XIST (spanning exons 5 and 6) were used for qRT-PCR analysis of XIST RNA in naïve (blue) and in vitro stimulated (orange) B cells. Relative fold change is shown, with values normalized to naïve B cells (set as 1). Standard error of the mean is shown with error bars.

**Supplemental Figure 2: macroH2A is not uniformly enriched on the Xi in activated B cells in healthy donors**. (A) Sequential XIST RNA FISH (red) followed by immunofluorescence detection of histone variant macroH2A (green). Representative field is shown, and arrowheads indicate macroH2A foci that co-localize with XIST RNA signal. (B) Quantification of XIST RNA localization patterns that co-localize with macroH2A foci. Number of nuclei counted is above each sample. (C) qRT-PCR analyses of macroH2A1.1 and macroH2A1.2 transcripts in naïve and in vitro stimulated (day 2) B cells (*left*). qRT-PCR analyses of both macroH2A1 variants for naïve and stimulated B cells compared to 293T, a human embryonic kidney fibroblast cell line (*right*). Statistical significance determined using one-way ANOVA across three cell types for each variant.

**Supplemental Figure 3: Sequential XIST RNA FISH and IF detection of the active chromatin modification H3K4me3**. (A) Representative field image for XIST RNA FISH (red) and sequential IF for H3K4me3 (green). The arrowheads indicate H3K4me3 ‘holes’ that overlap XIST RNA signals. (B) Quantification of XIST RNA localization patterns and H3K4me3 ‘holes’. Number of nuclei counted is above each sample.

**Supplemental Figure 4: XIST RNA localization patterns for naïve and *in vitro* stimulated B cells (day 2) from pediatric SLE patients and healthy age-matched controls**. (A) Quantification of XIST RNA localization patterns for naïve B cells. Number of nuclei counted is above each sample. (B) Quantification of XIST RNA localization patterns for *in vitro* stimulated (day 2) B cells using CpG. Number of nuclei counted is above each sample. (C) Representative flow cytometry analysis for CD86 staining, using unstained cells, naïve B (unstimulated), and day 2 stimulated B cells for a healthy control sample and one pediatric SLE patient sample (SLE 17). (D) Quantification of XIST RNA localization patterns for *in vitro* stimulated memory B cells from pediatric SLE patients and healthy controls. Number of nuclei counted is above each sample. Statistical significance determined using two-tailed test with unequal variance, comparing SLE to healthy controls. P values for each localization pattern of XIST RNA are shown.

**Supplemental Figure 5: Flow cytometry analyses of B cell populations from PBMCs of adult and pediatric SLE patients and age-matched healthy controls**. (A) Typical gating strategy for B cell subsets from patient PBMCs recovered post-thaw, following B cell isolation and selection for CD27-cells. (B) CD27-B cell population percentages for healthy control (HC) and SLE patients. Average percentages are shown in bold, at the bottom of each group. Pediatric samples are denoted as “P2”. Statistical significance determined using two-tailed t-tests comparing healthy controls to SLE, for each B cell subset. Only ABCs were significantly different among SLE and HC samples, yet comprise less than 2% of total B cells.

**Supplemental Figure 6: XIST RNA localization patterns for *in vitro* stimulated B cells (day 2) from adult SLE patients and healthy age-matched controls**. Number of nuclei counted is above each sample. Statistical significance was determined using two-tailed unpaired t-test for each pattern of XIST RNA localization, comparing SLE to healthy controls. SLE samples 580-253 and 342-184 are male individuals, and are included as negative controls.

**Supplemental Figure 7: Correlations between XIST RNA localization patterns and autoimmunity comorbidities and medications**. Bar graphs show percent of activated B cells with XIST RNA localization patterns Type I-IV (A-D) in SLE patients with or without thyroid disease or Sjogren’s Syndrome. Red, green, and blue bar graph pairs represent significant differences between group means. Bar graphs show percent of activated B cells with XIST RNA localization patterns Type I-IV (A-D) for adult SLE patients. The paired bar graphs consist of patients taking (left) or not taking (right) the designated medication. Blue bar graph pairs represent significant differences between group means; statistical significance determined without correction for multiple comparisons with alpha=0.05. Each row was analyzed individually, without assuming a consistent SD. HCQ: hydroxychloroquine; mycophen. mofetil: mycophenolate mofetil. (E) SLE patients taking HCQ have significantly higher percentages of Type I XIST RNA localization patterns. Dotplot showing percentages of B cells for groups of HC and SLE patients for each type of XIST RNA localization pattern. Black circles represent individuals not taking HCQ; aqua triangles are SLE patients taking HCQ. Horizontal bars show median values of patients taking HCQ (aqua) or other medications (black). Only Type I XIST RNA patterns were statistically significant among SLE patients for HCQ treatment. Statistical significance determined using two-tailed t test.

**Supplemental Figure 8: Linear regression analyses of SLE patient disease parameters**. (A) SLE disease parameters (ANA Titer, SLEDAI score, disease duration), patient age, and sample draw date were correlated with each XIST RNA localization pattern for *in vitro* activated adult SLE B cell samples. (B) Correlation between age and XIST RNA localization patterns for SLE patients (*left*) and healthy controls (*right*). XIST RNA Type IV patterns increased with age for SLE patients; there were no significant correlations between age and XIST RNA localization patterns for healthy controls.

## Acknowledgements

We would like to thank C. Berry for statistical consultation for correlation analyses; N. Jiwrajka and L. King for assistance with editing of the manuscript, and all members of the Anguera lab for helpful discussions. This research was supported by a University Research Foundation grant, a American Chemical Society grant, a McCabe Foundation grant), NIH R21 AI124084, NICHD 5K12 HD085848-03, NIH R01 AI 134834, DOD grant LR170055: W81XWH-18-1-06 (to MCA); and NIH 1F32AI154797 to SP.

## Author contributions

S. Pyfrom performed the bioinformatic analyses for single-cell RNAseq data sets, the bioinformatic analyses using human activated B cell RNAseq data sets, and the linear regression analyses for XIST RNA localization and comorbidities, and made corresponding supplementary tables and figures. B. Paneru performed the human B cell isolation and *in vitro* culture, the XIST RNA FISH and IF experiments on human B cell subsets, and quantified the localization patterns for SLE patients and controls. J. Knoxx and B. Paneru performed the flow cytometry and FACS isolation experiments. MCA made figures 1, 2, 3, 5, 6 in main text and supplementary figures S1, S2, S3, S4, S5, S6, S6. MCA and S. Pyfrom wrote the manuscript.

