## supplemental figures for "The dynamic epigenetic regulation of the inactive X chromosome in healthy human B cells is dysregulated in lupus patients"

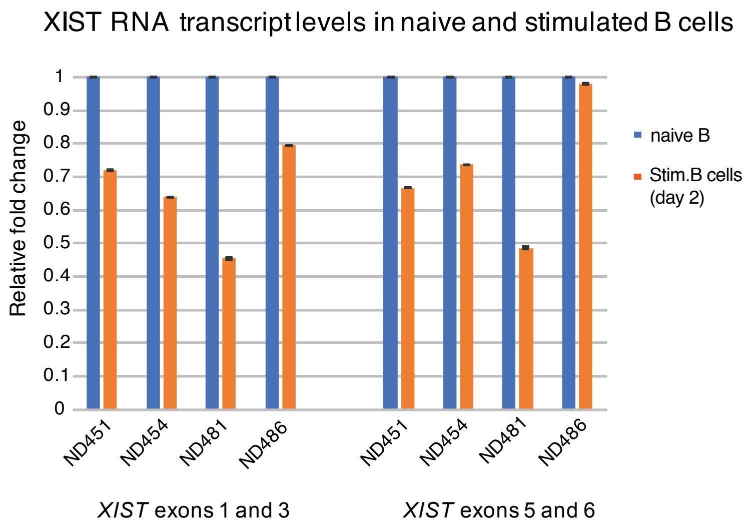

Supplemental Figure 1

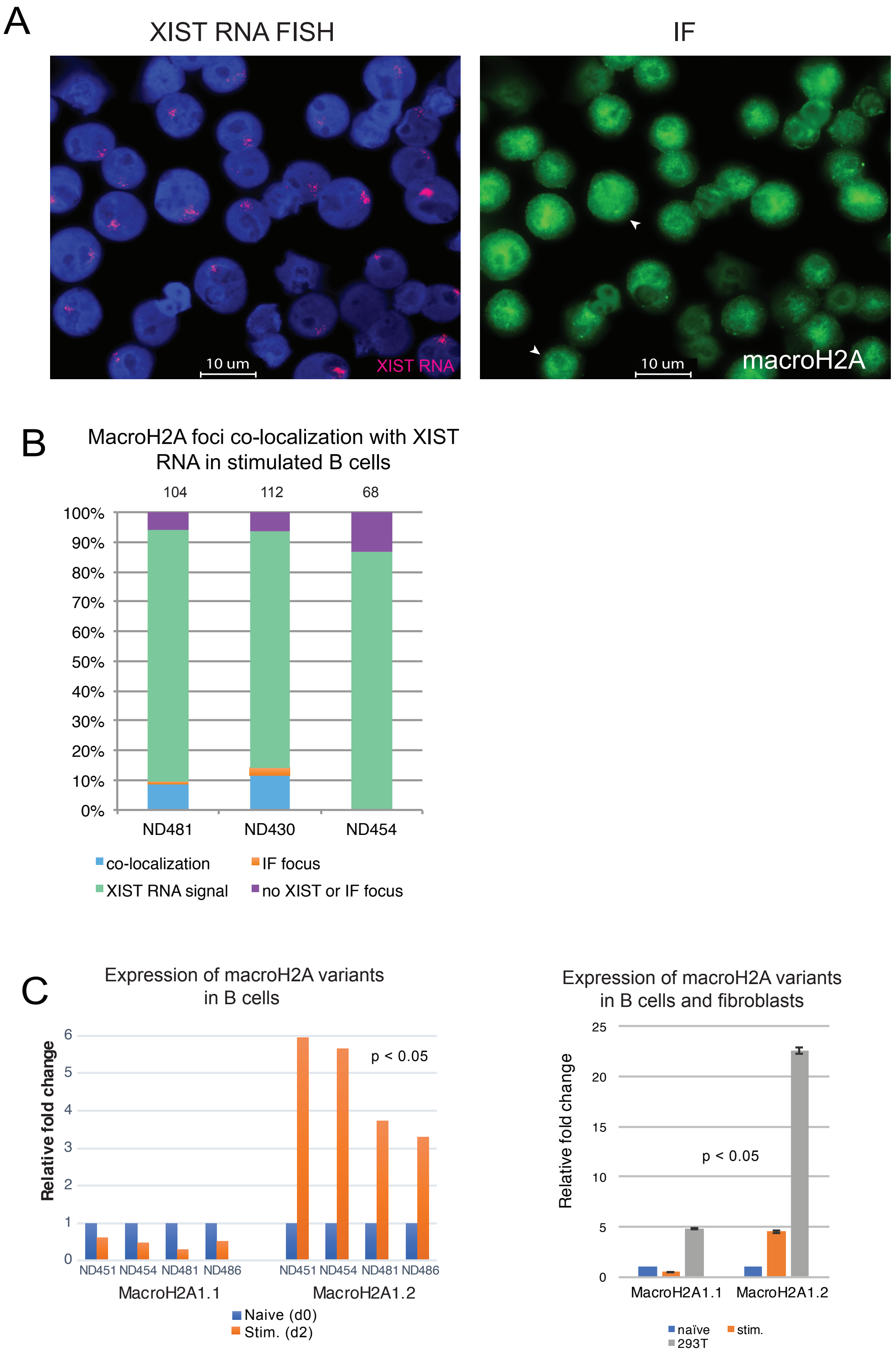

Supplemental Figure 2

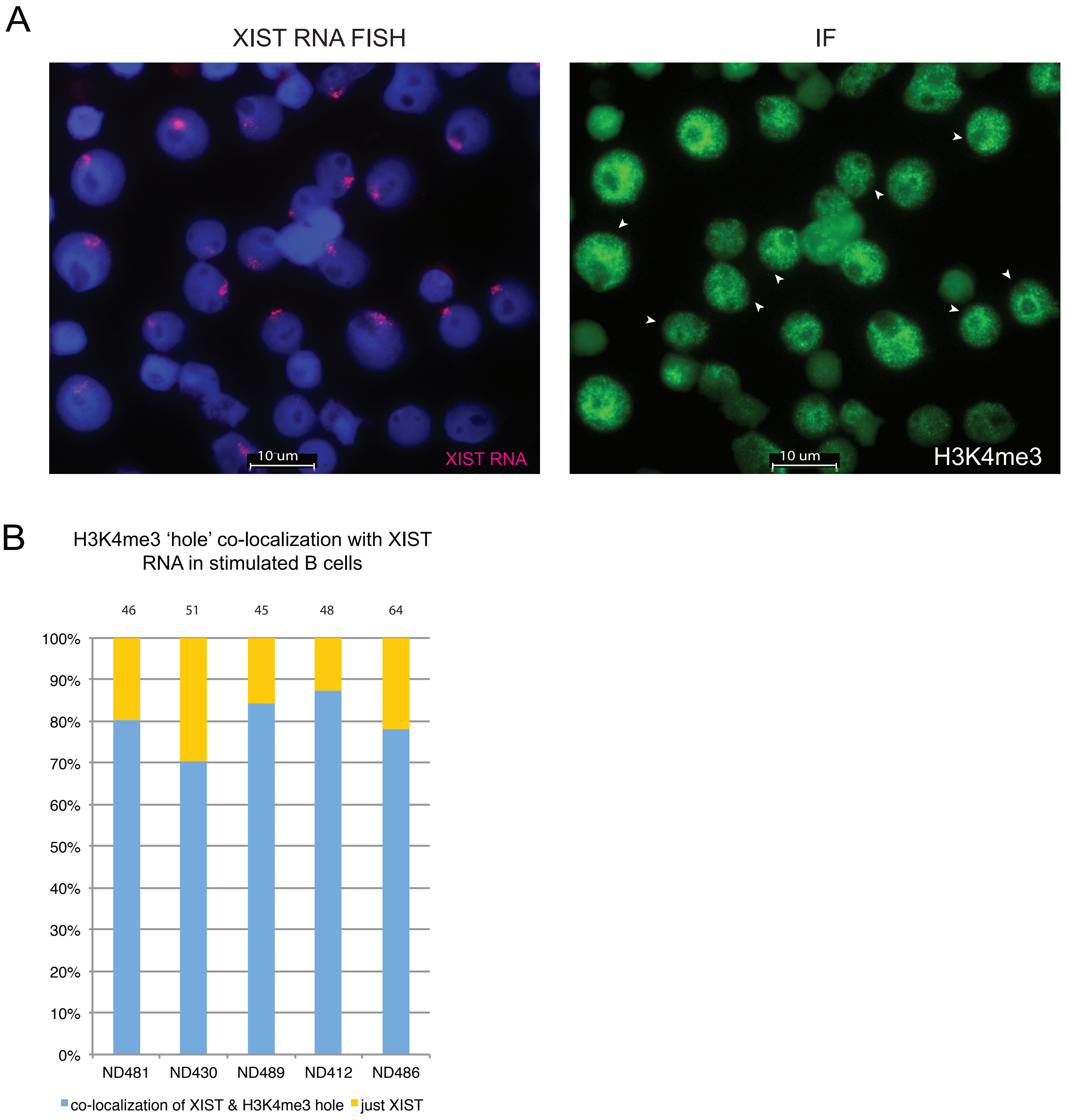

Supplemental Figure 3

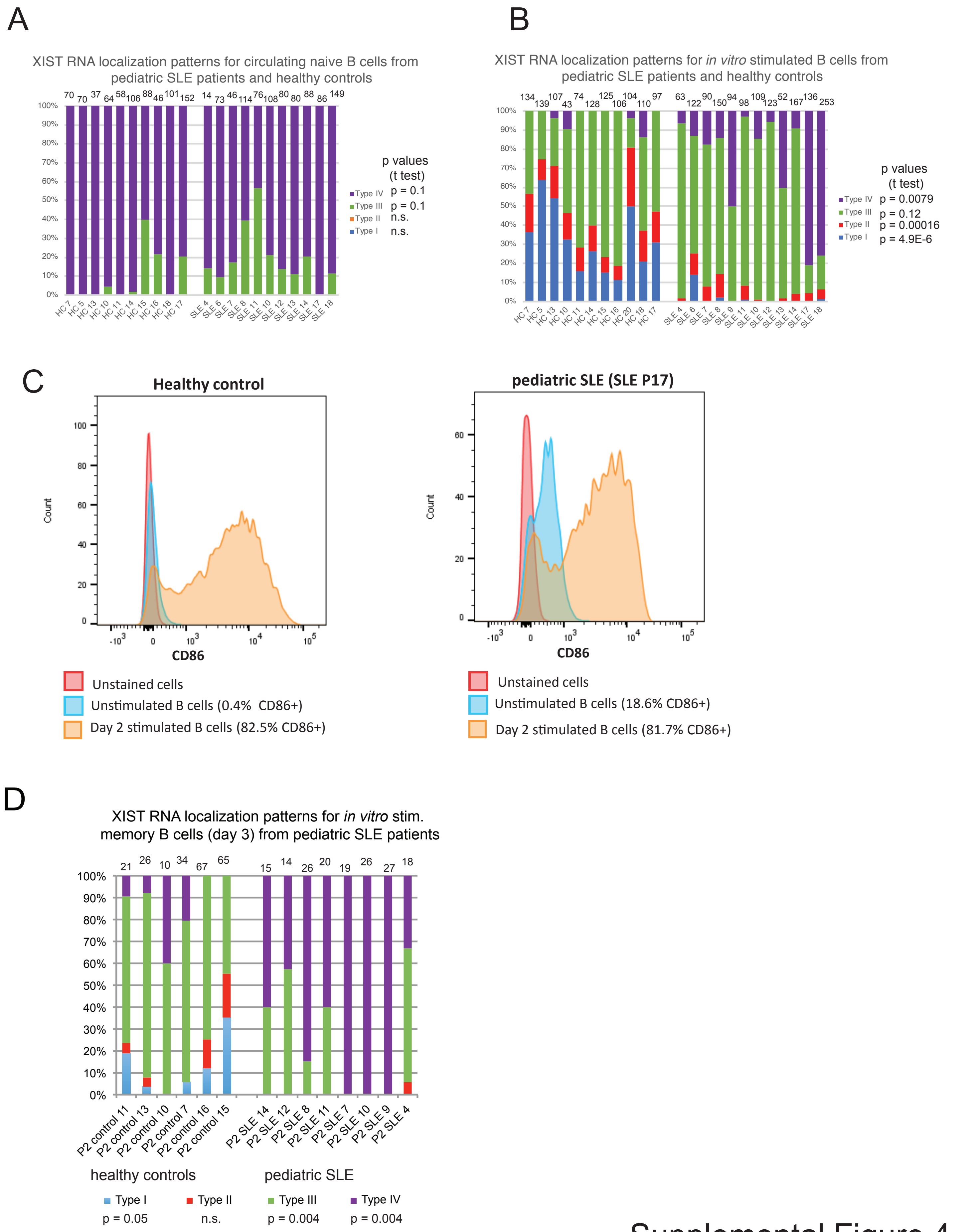

Supplemental Figure 4

A

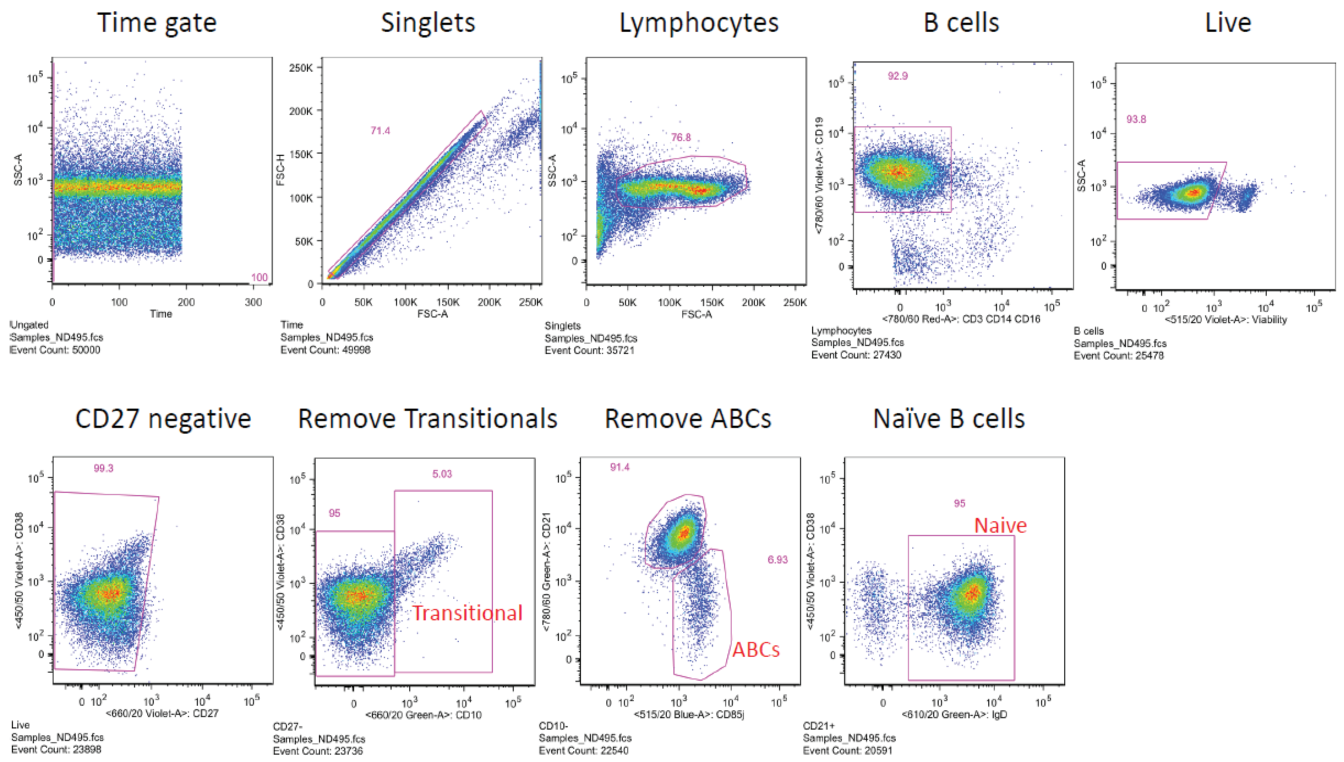

B

| Donor ID | Disease status | % lymphocytes | %B cells (of lymphocytes) | %live B cells | %CD27- | %Transitional B (CD10+CD38+) | %ABCs (CD10-CD21-CD85j+) | %Naive B (CD10-CD21+IgD+) |
|---|---|---|---|---|---|---|---|---|
| P2 control 18 | HC | 31.2 | 91.2 | 99.3 | 98.5 | 9.32 | 0.966 | 84.2 |
| Control 837 | HC | 42.2 | 96.6 | 98.1 | 99.2 | 7.45 | 1.04 | 83 |
| Control 601 | HC | 13.9 | 94.5 | 97.8 | 98.7 | 11.4 | 1.85 | 78 |
| 520-413 | HC | 54.9 | 83.1 | 95.2 | 97.3 | 11.5 | 1.03 | 77.6 |
| 458-188 | HC | 40.6 | 84.9 | 96.3 | 98.7 | 13.4 | 2.6 | 76.4 |
| 831-592 | HC | 35 | 77.1 | 95.6 | 97.2 | 24.4 | 1.24 | 62 |
| 434-601 | HC | 30.2 | 71.9 | 97.2 | 98.5 | 9.73 | 0.243 | 82.7 |
| 184-906 | HC | 58.8 | 63.1 | 78 | 99.6 | 14.8 | 0.293 | 78.8 |
| 206-018 | HC | 63.5 | 80.1 | 95.2 | 98.5 | 13.2 | 0.141 | 81.5 |
| 209-880 | HC | 11.8 | 68.5 | 77.7 | 99.7 | 11 | 1.13 | 77.5 |
| 225-065 | HC | 78.8 | 87 | 73.2 | 99.4 | 10.4 | 1.22 | 81.3 |
| 262-165 | HC | 36.4 | 75.6 | 72.4 | 99.5 | 7.03 | 1.41 | 78 |
| 306-826 | HC | 27.6 | 68 | 79.4 | 99.7 | 11.4 | 1.73 | 78.2 |
| 259-535 | HC | 31.8 | 79.8 | 85.8 | 97.4 | 19.4 | 1.74 | 68 |
| **Average** | | | | | | **12.45928571** | **1.188071429** | **77.65714286** |
| P2lupus17 | SLE | 73.1 | 95.9 | 98.1 | 99.5 | 4.09 | 1.55 | 87.5 |
| P2 lupus 18 | SLE | 70.4 | 94.3 | 89.5 | 99.5 | 16.8 | 1.31 | 77.4 |
| 271-643 | SLE | 51.3 | 79.9 | 98.1 | 99.4 | 10.9 | 0.428 | 83.1 |
| 280-535 | SLE | 59.2 | 86.5 | 96.7 | 98.8 | 13.7 | 2.91 | 77 |
| 100-221 | SLE | 50.3 | 88.3 | 96.3 | 97.9 | 11.4 | 2.12 | 78.8 |
| 567-547 | SLE | 47.1 | 83.5 | 97.1 | 99.3 | 15.6 | 3.13 | 69.9 |
| 106-930 | SLE | 65 | 64.3 | 58.9 | 99.6 | 24.6 | 0.182 | 71 |
| 580-253 | SLE | 56.5 | 65.5 | 80.7 | 99.9 | 8.32 | 0.73 | 86.3 |
| 126-590 | SLE | 63.1 | 71.5 | 80.3 | 98.1 | 2.41 | 1.44 | 88.4 |
| 142-961 | SLE | 60.8 | 67.8 | 73.9 | 99.8 | 26.1 | 3.96 | 66.3 |
| 147-591 | SLE | 64 | 76.4 | 75.2 | 99.5 | 3.14 | 3.14 | 84.8 |
| 150-646 | SLE | 60.3 | 58.4 | 22.9 | 97.9 | 3.83 | 0.851 | 86.8 |
| 211-152 | SLE | 74.9 | 84.3 | 69.4 | 99.4 | 4.75 | 1.74 | 83.5 |
| 212-918 | SLE | 69.9 | 77.8 | 79.9 | 99.5 | 1.3 | 0.518 | 92.5 |
| 304-618 | SLE | 52.6 | 78.1 | 94.5 | 98.4 | 2.21 | 1.25 | 89.2 |
| 335-682 | SLE | 81.8 | 79.1 | 65 | 99.3 | 21.6 | 1.61 | 73.3 |
| 341-163 | SLE | 49.4 | 73.8 | 81.7 | 98.7 | 11.2 | 2.39 | 78.6 |
| 400-053 | SLE | 67.9 | 76.5 | 83 | 98 | 36.8 | 1.9 | 55.3 |
| 411-496 | SLE | 58.7 | 77.2 | 76.2 | 99.6 | 6.83 | 2.6 | 77.7 |
| 422-812 | SLE | 42.9 | 73.2 | 65.9 | 99.6 | 4.14 | 1.53 | 88.9 |
| 559-114 | SLE | 54.8 | 68.8 | 85.4 | 99 | 33.3 | 3.86 | 54.5 |
| 666-788 | SLE | 48.1 | 73 | 90.2 | 98.7 | 2.34 | 0.937 | 91.4 |
| 670-247 | SLE | 39.1 | 67.6 | 62.4 | 98.7 | 0.512 | 3.32 | 83.6 |
| 750-330 | SLE | 66.7 | 69.9 | 53.8 | 99.9 | 13.6 | 2 | 76.6 |
| **Average** | | | | | | **11.64466667** | **1.891916667** | **79.26666667** |

two-tailed t-tests comparing SLE to HC:

Transitional B: n.s.

ABCs: $p = 0.035$

Naive B: n.s.

Supplemental Figure 5

A

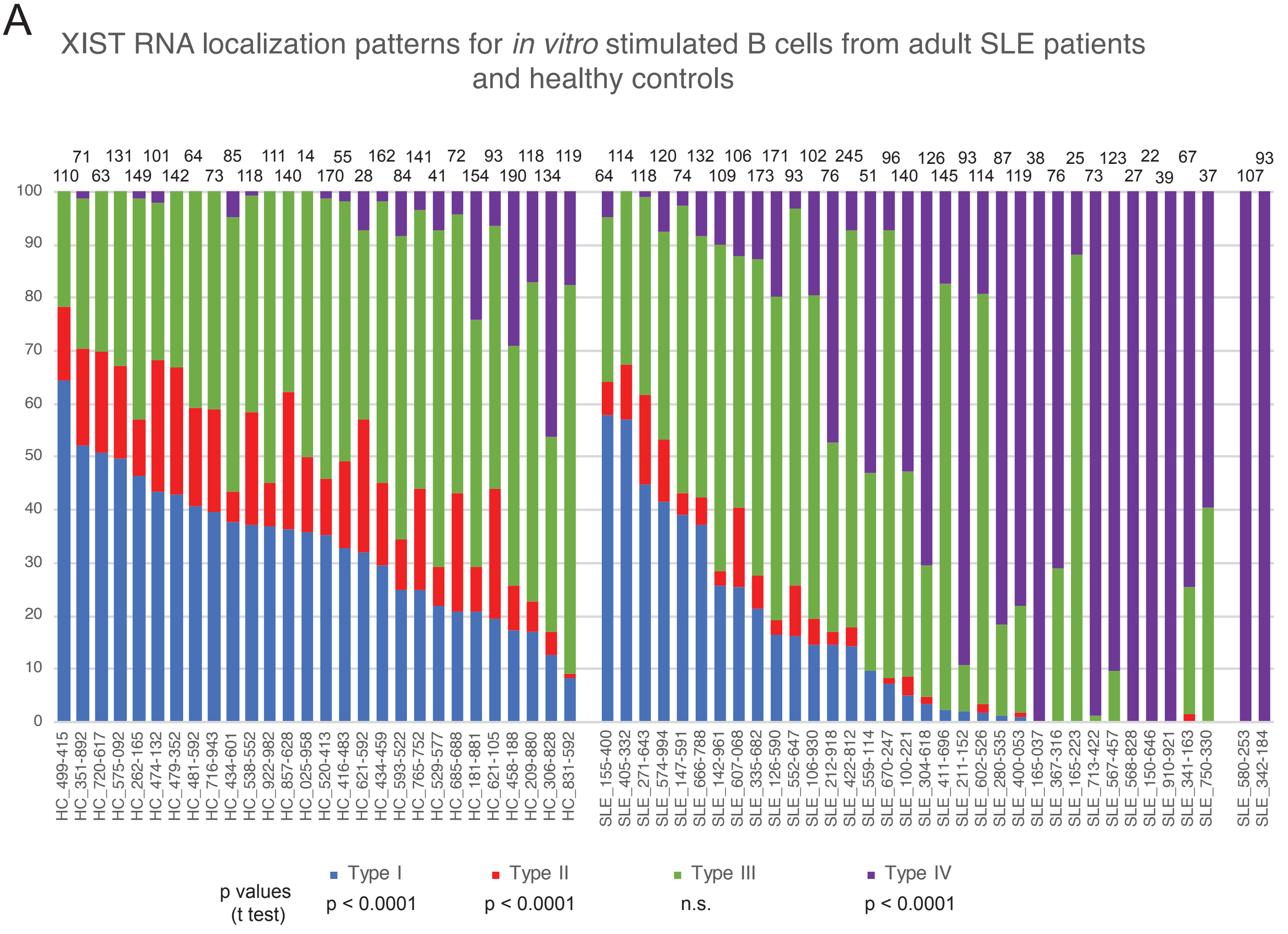

B

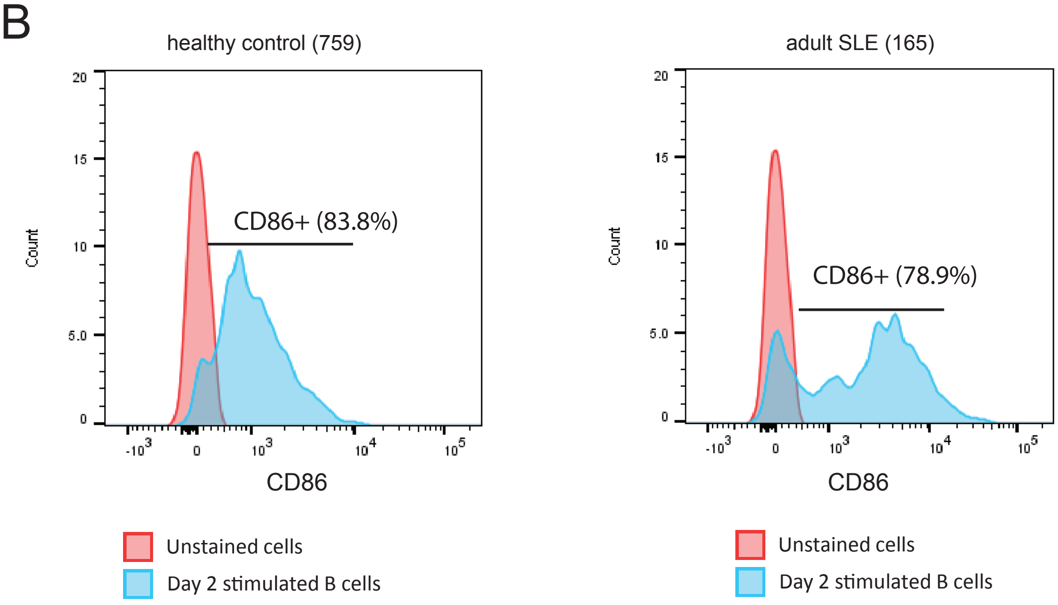

Supplemental Figure 6

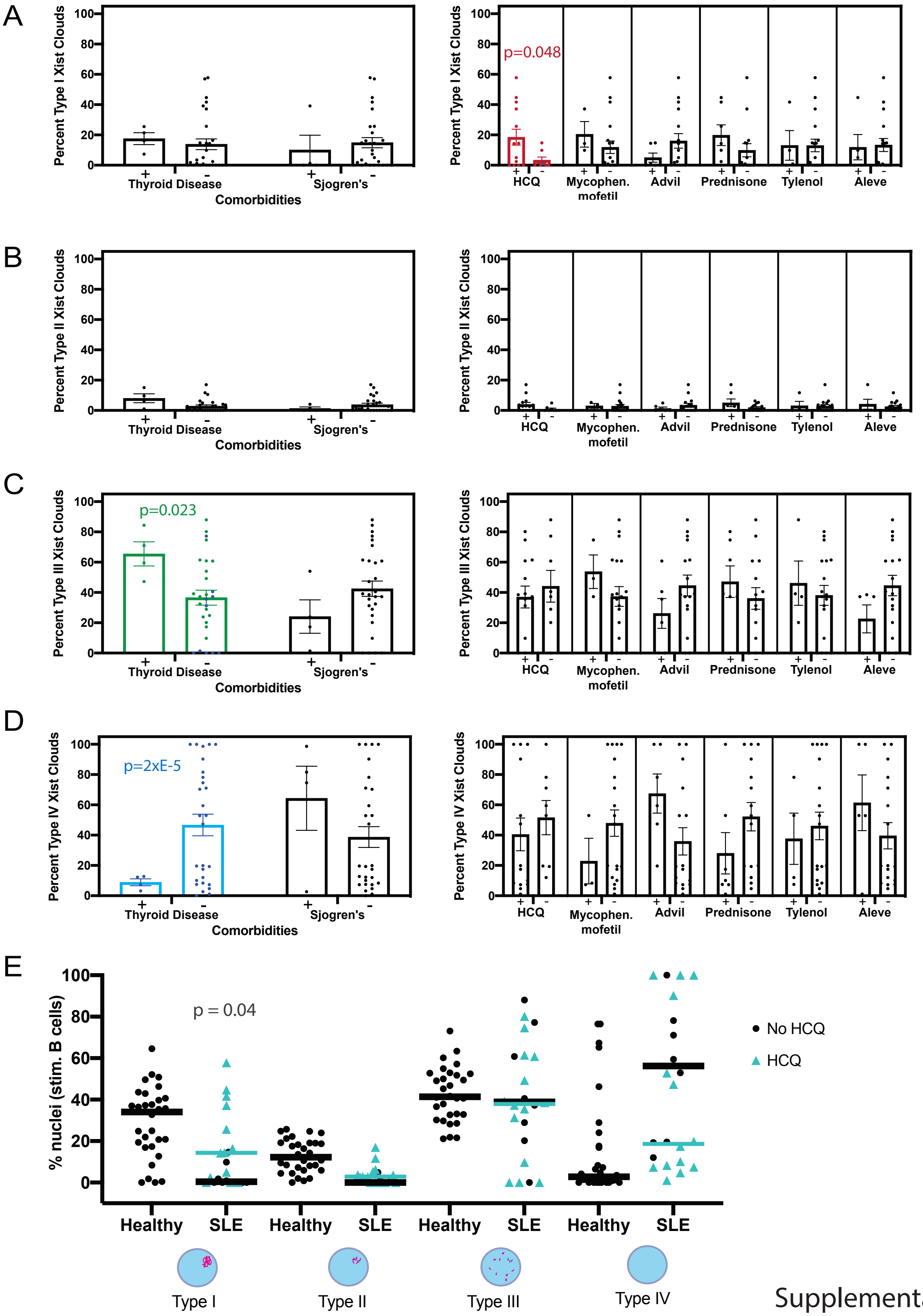

Supplemental Figure 7

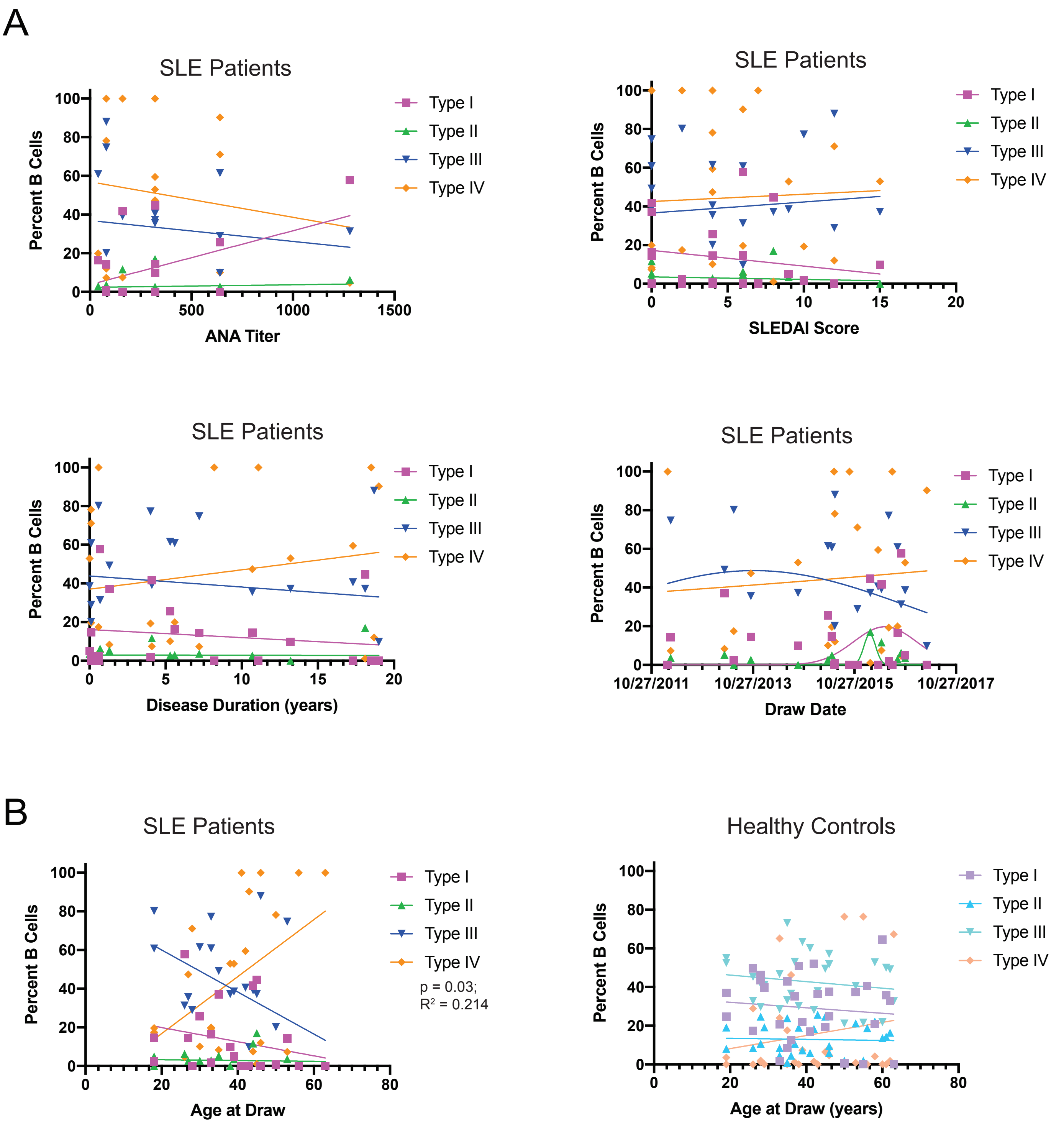

Supplemental Figure 8
